# Development of a multiplexed qPCRs-based approach for the diagnosis of *Dirofilaria immitis, D. repens*, *Acanthocheilonema reconditum* and the others filariosis

**DOI:** 10.1101/842575

**Authors:** Younes Laidoudi, Bernard Davoust, Marie Varloud, El Hadji Amadou Niang, Florence Fenollar, Oleg Mediannikov

## Abstract

**Background:** The frequent canine filariosis are caused by zoonotic filarial nematodes called *Dirofilaria immitis*, *D. repens* and *Acanthocheilonema reconditum* (Spirurida: *Onchocercidae*). The absence of reliable diagnostic tools to diagnose and discriminate between these infections as well as their different forms constitutes a major obstacle to their control. The serological diagnosis of heartworm disease has recently shown both sensitivity and specificity problems.

**Findings:** Herein, we developed and set up a novel molecular approach for an improved detection of the occult and non-occult filarioses, especially those caused by *A. reconditum*, *D. immitis* and *D. repens* as well as their differential diagnosis based on qPCRs assays. This approach designated the “Combined multiplex approach”, proceeds as follows: Filaria and wolbachia identification using the newly customized 28S-based pan-filarial and 16S-based pan-wolbachia qPCRs, respectively, followed by the fast typing method of positive samples using the triplex qPCR targeting *A. reconditum, D. immitis* and *D. repens*, and a duplex qPCR targeting *Wolbachia* of *D. immitis* and that of *D. repens*. The analytical sensitivity of the newly qPCR systems was confirmed by the detection limit of wolbachia and filaria DNA ranged from 5E^-1^ to 1.5E^-4^ mf/ml of blood with an R² higher than 0.99, Cohen’s Kappa agreement ranged from 0.98 to 1. The approach was complemented by a pan-filarial COI and pan-*Wolbachia fts*Z PCR for the identification of other filarial parasites and their *Wolbachia*, respectively.

When tested on clinical samples, the results are as follows: 29.2 % (49/168) tested positive to at least filariae or wolbachiae DNA. 19 samples of them tested positive for filarial DNA, 9 for wolbachia DNA and 21 for both. Filarial species and *Wolbachia* genotype were also identified by the combined multiplex approach from all the positive samples. The single DNA of *D. immitis* was identified in 12 samples, *D. repens* in 7, and *A. reconditum* in 15 samples, the co-infection was observed in 5 samples, 4 for both *Dirofilaria* and one harbored the three species. Therefore, 22 samples were positive for *Wolbachia* endosymbiont of *D. immitis*, 3 for that of *D. repens* and 5 for both genotypes. A newly duplex qPCR developed for the differential diagnosis of heartworm and French heartworm (*Angiostrongylus vasorum*) was successfully validated *in vitro*. However, no DNA of this latter was detected in canine blood samples used in this study. The immunochromatographic test for dirofilariasis antigen during evaluation before and after thermal pretreatment of sera showed substantial agreement (K=0.6) and weak agreement (K=0.15), respectively.

**Conclusion:** The proposed molecular tool targeting filarial genes and associated *Wolbachia* genes is a reliable tool for the exploration and diagnosis of occult and non-occult canine filariasis. We believe that the current diagnosis of heartworm based on antigen detection should be always confirmed by qPCR-based essays; the heat-pretreatment of sera is useless and strongly discouraged.

## Background

Canine filariosis include diseases caused by parasitic nematodes called filaria, belonging to the order Spirurida. There are several species of veterinary and human importance. Dogs seem to be the natural hosts for several species, such as *Dirofilaria immitis, D. repens, Acanthocheilonema reconditum, A. dracunculoides, Cercopithifilaria grassii, Brugia ceylonensis, B. patei, B. malayi, B. pahangi, Onchocerca lupi* and *Thelazia callipaeda* [1, 2, 3, 4]. These arthropod-borne filarioids produce blood, cutaneous or mucous microfilariae, where they are available for arthropod vectors [5]. The most common and medically important species affecting dogs, are *D. immitis,D. repens* and *A. reconditum* [6]. In addition to their veterinary importance, they can also affect human health, where *D. immitis* was recently considered as an emerging disease in Europe and causes the cardiopulmonary dirofilariasis. In the USA, it is the most important parasitic infection threatening dog’s life [7]. Elsewhere in the world, particularly in Eastern European countries, *D. repens* is the most endemic parasitic nematode causing subcutaneous infection, which is less virulent but more zoonotic than *D. immitis* [8]. Hence, both are emerging zoonotic diseases, whereas *A. reconditum* is an occasional zoonotic agent, which affects the subcutaneous tissue and the perirenal fat [9, 10] and causes the common infection in dogs, but is clinically less important [6].

Once became mature, these filarioids can produce blood circulating microfilariae. This larval stage (L1) is also a target for the diagnosis made by microscopic detection of these larvae or their DNA in the host blood [11]. *D. immitis,* the agent of heartworm disease, is distributed worldwide and is responsible for the heart failure in dogs after the colonization of pulmonary arteries and the right ventricle, where it can be fatal if untreated. Due to the gravity of the disease, it remains the most commonly diagnosed filariasis in dogs due to the detection of antigen circulating in the blood [11,12,13].

Several diagnostic method problems were raised, such as morphological confusion between the microfilariae of *D. immitis*, *A. reconditum* and *D. repens.* Commercially available diagnostic kits for the detection of *D. immitis* antigens may also cross-react with filarial and non-filarial nematodes, such as *D. repens*, *A. reconditum*, and *Onchocerca* spp. [15,16, 17], *Spirocerca lupi* and *Angiostrongylus vasorum,* especially the latter which can cause cross reaction without prior heat pretreatment of the sera [16, 17]*. A. vasorum,* the agent of French heartworm disease, should also be taken into account in the differential diagnosis of pulmonary disease [20]. So-called occult heartworm characterized by the absence of microfilaremia or an amicrofilaremia, this may result from the host’s immune response, low parasite load and infertility or, incidentally, the microfilaricidal effect observed in dogs receiving macrocyclic lactone prevention [17].When occult heartworm occurs in a co-infection with another filariosis, the diagnosis is even more challenging. The association of the heartworm with *D. repens* infection may result in an unexplained suppressive effect on the production of microfilariae of *D. immitis* [21]. In such cases, the cross reactivity between *D. immitis* and *D. repens* may result in misdiagnosis. Therefore, there is an urgent need for a reliable diagnostic method to detect occult as well as non-occult canine filariosis and to identify the pathogen. *D. immitis* detection has gained more and more attention; many trials have been performed for improving the quality of heartworm diagnostic tools, such as the detection of a specific antigen released by these worms [22], or the use of a recombined antigen of *D. immitis* for the specific antibodies detection [23].

The endosymbiotic intracellular bacteria of *Wolbachia* genus are associated with some filarial species of two subfamilies of *Onchocercidae: Onchocercinae* and *Dirofilariinae* [24]. These bacteria are host-specific, and each species of filarial worm is associated with a specific bacterial genotype. wolbachiae were targeted for the indirect diagnosis of *D. immitis* infection when occurred in the dead-end host like humans and cats, where the strong reaction of the host against parasites prevents them to achieve their maturation, and, therefore the production of microfilaria may not be achieved [24]. In such cases, the detection of filaria-specific *Wolbachia* may indicate a filarial infection and can serve as an alternative diagnostic tool in endemic areas [24, 25]. In around 40% to 60% of canine heartworm cases, both *Wolbachia* and parasite DNA may be detected using conventional PCR [26, 27, 33]. Indeed, the combined detection of *Wolbachia* and *Dirofilaria* DNA was suggested to improve heartworm detection [29].

In the present study, we developed a novel real-time PCRs-based assay for the rapid detection and identification of occult and non-occult canine filariosis and the French heartworm infection that can be used in routine diagnostic laboratory. We also evaluate the effectiveness of a new molecular approach to conventional serological diagnosis and evaluate the importance of serum heating.

## Methods

### DNA collection

Three groups of DNA were used to validate PCRs: (i) A total of 24 filarial and non-filarial nematode DNAs, collected in different countries (Table S1), were used to evaluate the specificity and sensitivity of designed oligonucleotide primers and probes; (ii) the DNA samples from eight strains of *Wolbachia* endosymbiont of *Aedes albopictus, Anopheles gambiae, Cimex lectularius, Cimex hemipterus* (PL13 strain), *D. immitis* microfilariae, *D. repens*, *Onchocerca lupi, Wuchereria bancrofti* and *Brugia* spp. were used additionally with the previous collection as control for customized designed qPCR targeting *Wolbachia* sp. and (iii) and 57 DNA from the frequents hosts and hemo-pathogens, that were considered negative controls, were used to confirm the primer’s specificity (Table S2).

### Blood and sera specimens

Canine blood samples were collected by a veterinarian using cephalic venipuncture into citrate and serum separator tube. The serum collected and the citrate blood were then stored at −20° C. This population is split into four groups: (i) 8 samples composed of nematode free laboratory Beagles from the biobank of veterinary research center of the IHU Mediterranee Infection were used as negative control group; (ii) 136 dogs enrolled on March 2017 from Corsica Island in which heartworms are endemic; (iii) 17 dogs from Côte d’Ivoire exhibiting blood microfilariae; and (iv) 7 military working dogs from France recruited on October 2018. Among them, 6 dogs were harboring microfilariae. Theses 168 samples were obtained and were subsequently processed for molecular and serological analysis.

### DNA extraction

The genomic DNA was extracted from the blood and the microfilaria-containing tissues using the QIAGEN DNA tissues extraction kit (QIAGEN, Hilden, Germany), following the manufacturer’s recommendations. The extracted DNA was eluted in a total volume of 100 µl and was then stored at – 30°C.

### Probes and primers sets design and run protocol

#### Custom protocol and *in silico* validation

After the choice of the target gene for each PCR, the sequences of the representative members of *Onchocercidae* family or *Wolbachia* genotypes were recovered from GenBank, then aligned using BioEdit v 7.0.5.3 software [30]. The highly conserved inter- and intra-species regions were used in Primer3 online software v. 0.4.0 (http://primer3.ut.ee), in order to determine a valuable candidate primers and probes taking into account the physicochemical characteristics, the default 60 and 65°C annealing temperature parameter were chosen for primers and probes respectively. Therefore, the *in silico* validation was made by a double check of each PCR system: (i) when the physicochemical characteristics, especially the hybridization temperature, hairpin, self and hetero-dimers formation, were checked using Oligo-Analyzer 3.1 [31] and (ii) primer-BLAST available at http://www.ncbi.nlm.nih.gov/tools/primer-blast [32], was used for checking system’s specificity in which all the expected or unexpected amplifications have been explored within the DNA databases of metazoans (taxid:33208), vertebrates (taxid:7742), bacteria (taxid:2), *Canidae* (taxid:9608), *Felidae* (taxid:9682) and humans (taxid:9605).

#### TaqMan simplex-qPCR targeting filarial nematodes

The choice of the large subunit rRNA gene (LSU), also called 28S gene, was based on several criteria such as: the tandem repetition of about 150 times in filarial nematode genome, which improves the PCR detectability [33], availability in GenBank for all families of nematodes and sharing a highly conserved region within *Onchocercidae* members. Primers qFil-28S-F, qFil-28S-R and a TaqMan® hydrolysis probe (qFil-28S-P) were designed to amplify most of filarial species (Table 1).

**Table 1.**
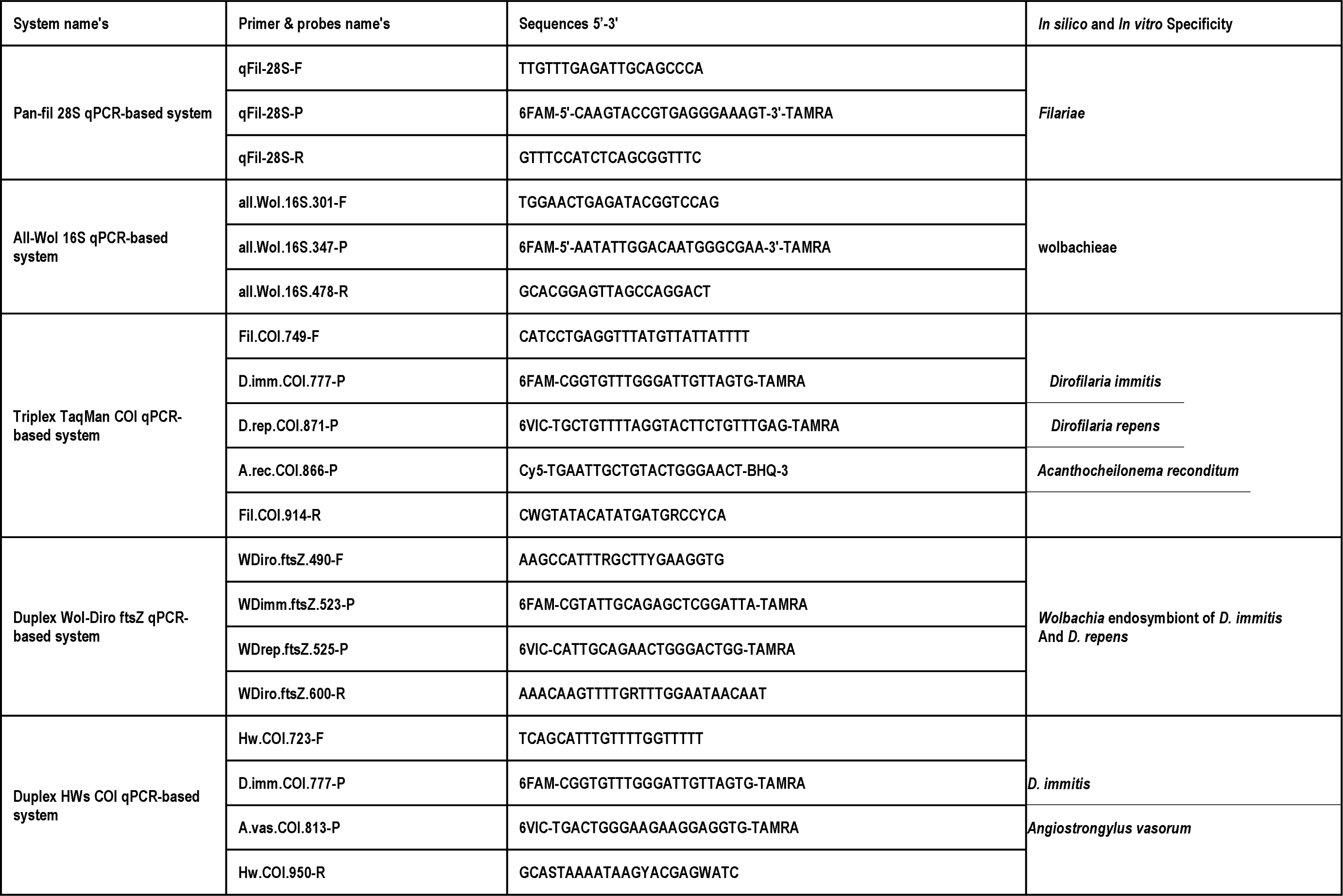
Primers and probes used in this study

#### TaqMan triplex-qPCR targeting *D. immitis, D. repens* and *A. reconditum*

The gene encoding for the cytochrome C oxidase subunit 1 (COI gene) was selected for the development of triplex TaqMan qPCR system targeting *D. immitis, D. repens* and *A. reconditum* (Table 1). This choice was based on the availability of COI for the three species in GenBank. The presence of this gene in several copies by the cells improves the sensitivity of the test. Indeed, COI has been described as a “barcode gene” for filarial nematodes [34]. Designed primers (Fil.COI.749 & dg.Fil.COI.914) target to amplify 166 bp-long fragments of COI of most *Onchocercidae* members (Table1). The system’s specificity was confined to the TaqMan probes, namely P.imm.COI.777 specific to *D. immitis*, P.rep.COI.871 specific to both *D. repens* and *Candidatus* Dirofilaria (Nochtiella) honkongensis affecting dogs and human in Japan [35]. Finally, the probe P.rec.COI.866 is specific to *A. reconditum.* In triplex TaqMan system, three different dyes were used for specific detection: FAM and VIC with a non-fluorescent quencher-TAMRA confined to *D. immitis* and *D. repens* probe’s respectively, Cyanine 5 (Cy5) with a non-fluorescent quencher-BHQ-3 for *A. reconditum* probe (Table 1).

#### TaqMan duplex-qPCR targeting heartworm and French heartworm

Duplex COI-based qPCR was designed (Table 1), namely Hw.COI.723-F and Hw.COI.950-R, were designed to amplify partial COI gene (227bp) of both filarial and non-filarial nematodes, including *D. immitis* and *A. vasorum.* The primers were chosen to be flanking the previously designed probe for *D. immitis*. In addition, we designed a new one specific for *A. vasorum.* The TaqMan probes were labelled with FAM and VIC respectively, with a non-fluorescent quencher TAMRA.

#### TaqMan simplex-qPCR targeting Wolbachiae

The rDNA gene 16 has been reported as the most commonly used gene for *Wolbachiae* phylogeny [36]. The simplex qPCR was developed and validated *in silico* from the conserved region of the first third of the 16S rDNA gene. The designed system (Table 1) is composed of primers named Wol.16S.301f and Wol.16S.478r with the probe Wol.16S.347p specifically targeting all members of the *Wolbachia* lineages.

#### TaqMan duplex-qPCR targeting *Wolbachia* sp. endosymbionts of filariae

The prokaryotic tubulin analogue protein (*ftsZ*) gene provides a sufficient discrimination between *Wolbachia* from supergroups C and D that found in filarial nematodes and those of supergroups A and D found in arthropods with higher divergence between filarial Wolbachiae of C and D supergroups [37].The *ftsZ*-based duplex qPCR was designed and consists of a set of primers named WDiro.ftsZ.490f and wDiro.ftsZ.600r targeting a filarial Wolbachiae belonging to the supergroup C, which includes those found in *Dirofilaria* sp. However, the specificity of the duplex was confined to probes wDimm.ftsZ.523p and wDrep.ftsZ.525p specific for *Wolbachia* of *D. immitis* and *D. repens* respectively (Table 1).

#### Run protocols

Simplex, duplex and triplex qPCR reactions were carried out in a final volume of 20 µl, containing 5 µl of DNA template, 10 µl of Master Mix Roche (Eurogentec). The concentration of each primer per reaction was 0.5 µl for simplex qPCR and 0.75 µl for both duplex and triplex, 0.5 µl of both UDG and each probe. Finally, the volume was completed to 20 µl using ultra-purified water DNAse-RNAse free. The TaqMan cycling conditions included two hold steps at 50°C for 2 min followed by 15 min at 95°C, and 39 cycles of two steps each (95°C for 30 s and 60°C for 30 s). These reactions were performed in a thermal cycler CFX96 Touch detection system (Bio-Rad, Marnes-la-Coquette, France) after activating the readers of the dyes used in each qPCR systems. After running protocol, the accumulation of the relative fluorescence units (RFUs) was recorded during the extension step of qPCRs and used to set-up the cut-off value of each TaqMan system, according to the formula described [38]. Here, we fixed the tolerance value at 5% for all systems. The qPCR reaction was considered positive only if it showed a RFUs value higher than Cut Off. All these models were performed using the CFX Manager Software Version 3[38].

### Primers sets design, PCR and sequencing protocols

In order to complete the molecular identification of filaria and *Wolbachia* species, we designed two sets of degenerated primers (Table 2): (i) Fspec.COI.957f; Fspec.COI.1465r targeting 509 pb from COI gene of filarioids, and (ii) Wol.ftsZ.363.f; Wol.ftsZ.958.r targeting 595 pb of *ftsZ* gene of *Wolbachia* lineages that may associated with filarioids. All PCR reactions were carried out in a total volume of 50 µl, consisting of 25 µl of AmpliTaq Gold master mix, 18 µl of ultra-purified water DNAse-RNAse free, 1 µl of each primer and 5µl of DNA template. The thermal cycling conditions were: incubation step at 95°C for 15 minutes, 40 cycles of one minute at 95°C, 30s for the annealing at a different melting temperature for each PCR assays (Table 2), 45s for elongation time at 72°C followed by a final extension of five minutes at 72°C (Table 1). PCR amplification was performed in a Peltier PTC-200 model thermal cycler (MJ Research Inc., Watertown, MA, USA). DNA generated through PCR reaction was purified using filter plate Millipore NucleoFast 96 PCR kit following the manufacturer’s recommendations (Macherey Nagel, Düren, Germany). Purified DNA was amplified using BigDye® terminator v3.3 cycle sequencing kit DNA in line with the manufacturer’s instructions (Applied Biosystems, Foster City, USA). Sequencing was performed using 3500xL Genetic Analyzer (3500xL), the capillary electrophoresis fragment analyzer (Applied Biosystems®). The nucleotide sequences were assembled and edited using ChromasPro 2.0.0 and were then checked using Basic Local Alignment Search Tool (BLAST) [39].

**Table 2:**
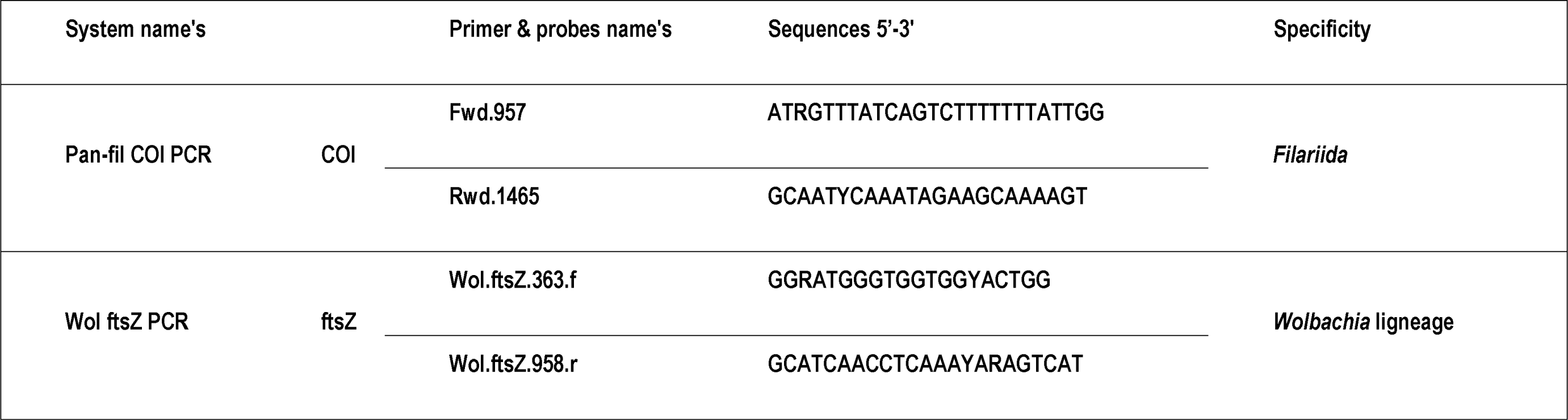
primers sets used for standard PCR and sequencing analysis.

### Specificity, sensitivity and *in vitro* validation of qPCR systems

In order to assess specificity of customized designed qPCR systems, we used our DNA collections described above. Furthermore, the specificity of multiplex qPCR systems was also assessed using individual and pooled DNA, where the DNA of *D. immitis, D. repens* and *A. reconditum* were used to validate the triplex, *D. immitis* and *A. vasorum* were used to validate duplex targeting heartworms and, finally, the wolbachiae DNA extracted from *D. immitis* and *D. repens* microfilariae were used as positive control to validate duplex system targeting these *Wolbachia*. The analytical sensitivity was assessed using a 10-fold dilution of DNA templates, and then standard curves and derived parameters (PCR efficiency, Slope, Y-intercept and correlation coefficient) were generated using CFX Manager Software Version 3 [38]. The triplex and pan-filarial qPCR systems were challenged in detecting the related microfilariae number of *D. immitis*, *D. repens* and *A. reconditum*. The DNA of each species was obtained from a naturally infected canine blood *D. immitis* (Corsica 2018), *D. repens* (Marseille 2018) and *A. reconditum* (Côte d’Ivoire, 2018). Firstly, 1 mL of each blood was examined by the modified Knott test [9] to identify the microfilaria species and their number, then the microfilaria concentration was ajusted at 1,500 mf/ml by adding Hank’s balanced salt solution (GIBCO®). Therefore, two extractions were performed from 200 µl, (i) from each separately calibrated sample and (ii) after mixing an equal volume of each of them to generate 500 mf/ml per ml of each species. They were used to evaluate the pan-filarial and triplex qPCRs respectively. Finally, a serial 10-fold dilution of DNA extracted from microfilaremic blood (Corsica 2018) containing 4,033 microfilariae of *D. immitis* was used to assess the analytical sensitivity of both triplex and duplex (Wol-Diro *ftsZ*) in the direct and the indirect detection of the single infection by *D. immitis*.

### Serological analysis

The serological analysis was performed using DiroChek® heartworm antigen test kit (Zoetis, Lyon, France) consisting of an enzyme-linked immunoabsorbent assay known as sandwich ELISA, targeting the antigen secreted by adult female heartworms [17]; where two antibodies directed against these antigens were used. The first one is coated in the wells and the second is labeled with the enzyme horseradish peroxidase, the revelation is performed after the addition of chromogenic substrate. However, the reading is subjective, where any change of color is considered positive. Each serum was tested using two different protocols: (i) in the first one, a volume of 200 µl of each serum was heated at 104°C for 10 minutes followed by centrifugation at 16,000 g. according to M.J. Beall’s recommendations regarding the immune complex dissociation to detect any heartworm antigen if present [40], (ii) the second protocol was performed without heat-treatment of the sera.

### Diagnostic approach standardization on biological samples

In addition to the heartworm antigen detection, we used the following approaches to explore and identify filarial infections from the biological samples described above. First, all samples were screened for both filaria and its Wolbachia DNA using the pan-filarial and pan-Wolbachiae qPCRs, respectively. Then, a partial COI and *ftsZ* genes were amplified and sequenced according to the previous protocol from all positive samples for filarioids and Wolbachia respectively. Secondly, the fast typing method based on the direct identification of filaria and their wolbachia using the approach combining the triplex COI and duplex Wol-Diro *ftsZ* qPCR based-systems. Finally, all samples were screened for heartworms using the Duplex HWs COI qPCR-based system in order to differentiate between *D. immitis* and *A. vasorum* DNA. In order to evaluate the performance of molecular and serological assays in the absence of gold standard test (necropsy), we developed the following approach to determine the true positive samples. The sample is considered truly positive for heartworm if the sample is positive for: 1) at least one of the molecular markers of heartworm (*D. immitis* DNA or that of its *Wolbachia*) and 2) antigen-test positive after immune complex dissociation by heating sera. This approach eliminates false-positive serological results that may be obtained by increasing the detection threshold (sensitivity) after heat-treatment of the sera before use, which is subsequently confirmed by the molecular markers specific to *D. immitis*. Once the DNA of *A. reconditum* identified, the sample is considered true positive, as well as for the DNA of *D. repens* or that of its *Wolbachia* which were considered as true positive samples for *D. repens*.

### Statistical analysis

Results generated through laboratory analysis were recorded in Microsoft Excel® program (Microsoft Corp., Redmont, USA). In order to assess how *Wolbachia* strains correlated with filarial species, the Spearman correlation coefficient was calculated using correlation/association tests. In order to evaluate the relevance of each diagnostic approach, the prevalence, correct classification, misclassification, sensitivity, specificity, false positive rate, false negative rate, positive, negative predictive value and Cohen’s Kappa (k) measures agreement were calculated. According to the scale of Landis and Koch (1977), the agreement quality of Kappa values was described as: < 0: no agreement, 0 - 0.2: small, 0.2 - 0.4: fair agreement, 0.4 - 0.6: moderate, 0.6 - 0.8: substantial, 0.8 – 1: almost perfect. The statistical analysis was performed using Addinsoft 2018(XLSTAT 2018: Data Analysis and Statistical Solution for Microsoft Excel, Paris, France).

## Results

### Systems validation

The *in silico* validations revealed that, the pan-filarial systems (28S qPCR and COI PCR) were specific for filarial parasites belonging to Dirofilariinae, Onchocercinea, Setariinae, Oswaldofilariinae, Icosiellinae and Waltonellinae subfamilies. The 16S qPCR targeting *Wolbachia* strains was specific for all the lineages known so far. However, the *ftsZ* PCR showed specificity for *Wolbachia* strains belonging to supergroups C, D, F and J, that may be associated with filarioids. Likewise, the multiplex qPCRs were also specific for the target species without failures. Additionally, *in silico* assessment of sensitivity was confirmed by the equality of melting temperature inside each set of primers. This temperature is less than that of probes. Indeed, the absence of primer-dimmer formation and hairpin structures was also confirmed. Furthermore, the specificity was confirmed again by an *in vitro* validation as shown in Table S1, where the positive reaction was obtained only from the target DNA and no negative control was amplified. Despite using single or pooled DNAs, the specific fluorescence signals generated through the multiplex qPCR systems were successfully related to the target DNA (Table S3, Fig.S1).

### Assays performance characteristics

The assay characteristics were assessed for the pan-filarial, the triplex and the duplex targeting *Wolbachia* endosymbiont of *Dirofilaria* spp. The analytical sensitivity of the pan-filarial qPCR was confirmed three times using *D. immitis, D. repens* and *A. reconditum* DNAs sharing the same microfilariae concentration. This assay was able to detect up to 1.5 ×10^-4^ mfs/ml (i.e., corresponding to 0.75 x 10^−6^ mfs/5μl). Efficiencies ranged from 99.3 to 104.3%, with a Slope from −3.34 to −3.22, Y-intercept values from 21.72 to 23.15 and an *R^2^* from 0.996 to 0.998 for all microfilaria species (Table S4 and Fig. S2). However, the analytical sensitivity of the triplex when using the pooled DNA of three species was confirmed by the detection of up to 5 ×10^-1^ mfs/ml (i.e., corresponding to 2.5 x 10^−3^ mfs/5μl) of each species simultaneously (Table S5, Fig. S3). qPCR efficiencies ranged from 100.4 to 103.7%, with a Slope from −3.23 to −3.3, Y-intercept values from 32.89 to 33.21 and an *R^2^* from 0.993 to 0.999. Finally, the analytical sensitivity was confirmed for both COI-triplex and *ftsZ* duplex qPCRs in detecting the single infection by *D. immitis* (Table S6).The detection limit was up to 5 ×10^-2^and5 ×10^-1^ mfs/ml, respectively (i.e., corresponding to 2.5 x 10^−(3 and 2, respectively)^ mfs/5μl), qPCR efficiencies were 104.8 and 100.5%, Slopes were −3.212 and −3.3, respectively, with an *R²* higher than 0.995 for both (Table S6, Fig. S4).

### Molecular diagnostic approaches

Results of molecular screening followed by the sequence taping approach are detailed in Fig 2 A-B and Table S. Of the 168 samples tested, 49 (29.17%) were positive to at least one DNA. All positive results were grouped in: (i) 19 blood samples positive only for filarioids DNA, (ii) 9 positive only for wolbachiae DNA and (iii) 21positive for both filaria and wolbachiae DNA. Although a partial COI and *ftsZ* amplicon was amplified from all positive samples for filaria and wolbachia, respectively, COI gene sequence-based identification allowed the identification of the causative agent of filariosis in 35 (87.5%) samples out of 40 samples amplified by PCR (Fig. 2A). We reported 12 (30%) cases of *D. immitis*, 7 (17.5%) cases of *D. repens*, 15 (37.5%) cases of *A. reconditum* and one showed both *D. repens* and *A. reconditum* DNA. Noteworthy, the amplicon sequences were obtained separately from this latter after serial dilution of blood before DNA extraction. However, *ftsZ* gene sequence-based identification allowed the identification of *Wolbachia* genotype in 25 (83.33%) out of 30 samples amplified by PCR Fig. 2B. 22 (73.33%) of *Wolbachia* strain having a close relatedness (accession number and % identity) to the strains identified in *D. immitis*, and 3 (10%) similar to that identified in *D. repens* (accession number and % identity). However, sequence-based identification failed to yield sequences from amplified DNA in 5 cases for each system, which corresponds to 12.5% and 16.7% of filaria and wolbachiae DNA, respectively. The Combined multiplex approach based on the triplex COI qPCR targeting filariae and the duplex qPCR targeting Wolbachiae allowed the detection of the 49 samples previously considered positive on filariae markers (Fig.3A and Table S8). The triplex COI qPCR identified the relative species from all the positive samples for filariae (40, 100%). Of these, thirty-four (34, 85%) samples had only one filariae DNA, where *D. immitis* was identified in 12 (30%) of them, *D. repens* in 7 (17.5%) and *A. reconditum* in 15 (37.5%). The number of samples with two filarial DNAs was 5 (12.5%), of which 4 were positive for *D. immitis* and *D. repens* and one for *D. repens* and *A. reconditum*, the triple positivity was obtained for one sample only (2.5%). The duplex Wol-Diro-*ftsZ* allowed the identification of wolbachia genotype from the total positive samples of wolbachiae DNA. Twenty-one samples (21, 70%) were positive for *Wolbachia* endosymbiont of *D. immitis,* three samples were found positive for *Wolbachia* endosymbiont of *D. repens* and five samples (16.67%) were positive for both strains. Among the 168 samples screened by duplex qPCR for heartworms agents (*D. immitis* and *A. vasorum*), 17 (10.12%) were positive for *D. immitis* DNA and no *A. vasorum* DNA was detected.

### Link between *Wolbachia* genotype and filarial species within the infected host

The results of the distribution of filarioids markers obtained by multiplex qPCRs are shown in Fig. 3A and Table S8. Interestingly, most samples positive for filarioids DNA were also positive for *Wolbachia* (21/40, 52.5%). Indeed, *Wolbachia* DNA was associated with at least one *Dirofilaria* spp. in 80% (20/25). The result of correlation analysis between *Wolbachia* strains and *Dirofilaria* species are shown in Table 3.75% (9/12) of samples positive for a single DNA of *D. immitis* were also positive for *Wolbachia* genotype known to be associated with this filarioid, which corresponds to a significant correlation (r=0.509, p<0.0001). In addition, there is a significant correlation (r=0.181, p<0.019) between the presence of a single DNA of *D. repens* and that of the *Wolbachia* strain commonly associated with this filarioid, as well as samples containing both *Wolbachia* strains (r=0.454, p<0.0001). The presence of both *D. immitis* and *D. repens* DNA is mostly associated with the strain of *Wolbachia* endosymbiont of *D. immitis* (r=0.244, p<0.002) and secondarily, with both *Wolbachia* strains together (r=0.175, p=0.023). On the other hand, 29.63% (8/27) of samples harboring *Wolbachia* endosymbiont of *D. immitis* were free of filarioid DNA (Fig. 3A). 25% (2/8) of samples positive for *Wolbachia* endosymbiont of *D. repen s*DNA were also free for filarioid DNA. No correlation was observed between *A. reconditum* and *Wolbachia* strains.

**Table 3:** Cross tabulation showing the molecular exploration and sequence typing of filaria and wolbachia from canine blood.

|  | At least one<br>Filarioid DNA | *A. reconditum* | Positive for both<br>*Dirofilaria* spp. | Single DNA of *D.*<br>*repens* | Single DNA of *D.*<br>*immitis* |
| --- | --- | --- | --- | --- | --- |
| At least one wolbachiae<br>genotype | **0.506**<br>(**<0.0001**) | 0.053<br>(0.49) | **0.284**<br>(**0.0002**) | **0.447**<br>(**< 0.0001**) | **0.565**<br>(**< 0.0001**) |
| Positive for both<br>*Wolbachia* genotypes | **0.313**<br>(**<0.0001**) | 0.059<br>(0.449) | **0.175**<br>(**0.023**) | **0.474**<br>(**< 0.0001**) | **0.174**<br>(**0.025**) |
| Single DNA of<br>*Wolbachia* of *D. immitis* | **0.478**<br>(**<0.0001**) | 0.093<br>(0.230) | **0.305**<br>(**<0.0001**) | **0.419**<br>(**<0.0001**) | **0.606**<br>(**< 0.0001**) |
| Single DNA of<br>*Wolbachia* of *D. repens* | **0.334**<br>(**<0.0001**) | 0.018<br>(0.82) | 0.125<br>(0.105) | **0.458**<br>(**<0.0001**) | 0.11<br>(0.154) |

Values in bold are significantly different from 0.

### Heartworm antigen detection and infectious status

Of the 168 dog sera tested for heartworm antigen, 16.67% (28) were positive before pretreatment of the sera and were grouped into three groups (Fig. 3B): (i) 9 (5.36%) were mono-infected by heartworm and were positive for both *D. immitis* and its *Wolbachia* DNA except one, which was positive for *D. immitis* DNA only. (ii) 10 (5.95%) were co-infected samples with at least one other filarioid in PCR, consisting of 8 (4.76%) bi-infected with *D. repens* and 2 (1.19%) harbored molecular markers of *D. immitis, D. repens* and *A. reconditum.* (iii) Finally, 9 (5.39%) were positive for filaria other than heartworm: 7 (4.17%) samples positive for *A. reconditum* DNA only, 1 (0.6%) positive for *Wolbachia* endosymbiont of *D. repens* and 1 (0.6%) positive for both *A. reconditum* DNA and *Wolbachia* endosymbiont of *D. repens*.

Once the heat pre-treatment of sera was performed, the rate of positive samples increases up to 71.43% (120). 27.98% (47) of samples, harboring at least one DNA markers of filarial parasites or their *Wolbachia,* were also detected. However, two samples (1.19%), one positive for *Wolbachia* endosymbiont of *D. immitis* by qPCR and the other positive for that of *D. repens* by qPCR, remained serologically negative. 43.46% (73) were filarioids markers free that were considered positive for unknowing antigens (Fig. 3B and Table S9). No positive results in the negative control group for both serological and molecular assays were obtained.

### Performance characteristics comparison of the diagnosis approaches

Once the true positive samples of each filariosis were determined, the diagnostic value was evaluated for each test in the specific detection of filariosis. The sequence taping approach combining the filariae and wolbachiae identification allowed the diagnosis of heartworm infection in 86.21% (25/29) of cases, which corresponds to 82.8% and 99.3 of specificity and sensitivity, respectively, and allowing an almost perfect agreement with the true positive rate (K=0.87). The approach combining the multiplexed qPCRs was more sensitive than the gold standard in this study, in which all true positive heartworm samples were detected with another one positive for *Wolbachia* endosymbiont of *D. immitis* but negative by antigen test (Table S9). This combined multiplex approach allowed 100% and 99.3% of sensitivity and specificity respectively, with an almost perfect agreement of 0.98 (Table S10). While the heartworm antigen detection performed prior heat pre-treatment of sera showed 65.5 and 93.3% of sensitivity and specificity respectively, thus corresponding to a moderate agreement (0.6). In addition, the heat pre-treatment of sera allowed the detection of 71.4% (120) samples including 47 harboring filariae and wolbachiae markers and 73 with unknowing antigens. The performance characteristics of this tool in detecting heartworm infection when the sera are heated were 100% of sensitivity and 34.5% of specificity with a far agreement (0.15) (Table S9). The combined multiplex approach was considered as a gold standard for the diagnosis of *D. repens* and *A. reconditum.* The sequence typing method was 100% specific and sensitive at 62.6 and 94.1% for the specific detection of *D. repens* and *A. reconditum,* respectively. A substantial (0.75) and an almost perfect (0.97) agreements with the gold standard test were observed for the detection of *D. repens* and *A. reconditum,* respectively.

## Discussion

### Systems validation and assays performance characteristics

In this study we aimed to develop and set up a new molecular approach for the correct diagnosis of canine filariosis and their occult and non-occult forms. Once the PCR systems developed and validated, we evaluated its performance and characteristics according to four approaches: (i) one combining the screening followed by sequence typing of Filariae and wolbachiae, (ii) an other combining multiplexing qPCRs fast typing method, including *A. reconditum*, *D. immitis* and *D. repens* with the specific *Wolbachia* genotypes of these last two species, (iii) a serological assay using a commercialized heartworm antigen test with and without heat pre-treatment of sera, and (iv) finally, a differential diagnosis of heartworm with French heartworm (*A. vasorum*) using a duplex qPCR targeting both species. The newly customized PCR systems have shown a specific detection of the targeted DNA for which they were designed. The pan-filarial 28S qPCR system is presented for the detection of filarial DNA form biological samples. It has been adapted for the detection of filarial parasites known so far, such as the subfamilies Dirofilariinae and Onchocercinea, whose members parasitize mammals, reptiles and birds, and those of the subfamily Setariinae, confined to a large mammal, and Oswaldofilariinae parasites reptiles and subfamilies Icosiellinae and Waltonellinae amphibians parasites [5]. The LSU (28S) gene targeted by this system is known for its conserved regions between the filarial species [41]. The second qPCR system was customized for the detection of DNA from all *Wolbachia* lineages. The system provided a useful exploration of the presence of wolbachiae DNA despite their lineages, which is related in return to the conserved region of the 16S gene [36]. Another qPCR system for wolbachiae targeting the 16S gene was proposed as a complementary diagnosis of the lymphatic filariasis caused by *Wuchereria bancrofti* from human blood [42].

Multiplexing qPCRs were not only specific, but also discriminatory towards targeted DNA without failure (Table S3).These futures are directly related to the choice of the genes targeted, which offers a sufficient discrimination between species, as is the case with COI gene representing a nematode barcode [34][43], which is used for the development of the triplex qPCR for *D. immitis, D. repens* and *A. reconditum* and a duplex targeting *D. immitis* and *A. vasorum* agents of heartworm. Nonetheless, the *ftsZ* gene is often used for the characterization of *Wolbachia* supergroups, including those associated with filarial nematodes [37], from which a duplex qPCR targets both *Wolbachia* genotypes associated with *D. immitis* and *D. repens*. A quantitative real time qPCR targeting the *ftsZ* gene of *Wolbachia* endosymbiont of *B. pahangi* has been described [44].

It is interesting to note that molecular diagnosis combining the detection of filariae and *wolbachiae* DNA is an improvement and a tool for evaluating treatment protocols targeting filariae and wolbachiae [44]. The analytical sensitivity of new customized qPCRs ranged from 99.3% to 107.6%, with a slope value of standard curves ranging from −3.34 to −3.15 and coefficients of determination (R^2^) higher than 0.99. These characteristics are directly derived from the design protocol, where the formation of heterodimers and hairpins inside and between primers and probes was avoided. Primer sets share a similar melting temperature lower than that of the probes, giving a better sensitivity to the qPCR reaction.

The sensitivity of the pan-filarial 28S qPCR system was much higher than the triplex qPCRs for the detection of *D. immitis*, *D. repens* and *A. reconditum* DNA, where the detection limit was 1.5 E^-04^ and 5 E^-02^ mf/ml of blood, respectively. This difference could be explained by the tandem repetition of about 150 times of the 28S in filarial nematode [33], and also by the difference in stability of genomic and mitochondrial DNA sequences during the extraction protocol used in this study, which may have contributed to the difference in sensitivity. Therefore, the sensitivity of the triplex in detecting single DNA of *D. immitis* was also higher than the duplex *ftsZ* qPCR in detecting its *Wolbachia* and was able to detect up to 4.02 E-3 and 4.03 E-1 mf/ml, respectively. Difference in sensitivity of simplex qPCR for the detection of the filarioid *W. bancrofti* and the *Wolbachia* DNA from infected blood was 1-fold difference in copy number per cell in favor of filarial DNA, and can be explained by the fact that the eukaryotic DNA is more stable than bacterial DNA during the extraction [42].

### Molecular diagnostic approaches

Here, we developed and assessed two molecular approaches in detecting and identifying canine filariosis. The first one combined the screening and sequence typing of both filarial and *Wolbachia* DNA. The genomic DNA was identified with an almost perfect specificity ranging from 99.3 to 100%. However, the sensitivity ranged from moderate (62.5%) to perfect (94.1%) regarding the presence or not of co-infection. The overlapping peaks corresponding to different nucleotides on electropherograms of sequenced samples suggest the co-infection [40, 4]. The second approach combines two multiplexed qPCR systems targeting *A. reconditum*, *D. immitis*, *D. repens* and *Wolbachia* genotypes associated with the latter two. All samples showed at least one molecular marker which was detected and identified with an almost perfect sensitivity and specificity using this approach. The tool was fast, simple to use, sensitive and highly specific in detecting occult and non-occult filariosis within the infected hosts. This result reinforcing the utility of multiplex qPCR in detecting co-infections, confirms the limit of sequence typing method in the identification of co-infection [46], and avoids the classical sequence typing method [47].

### Linkage between *Wolbachia* strains and filarial species within the infected host

As expected, the *Wolbachia* DNA was significantly associated with *Dirofilaria* species in 80% (20/25), reinforcing the idea that this endosymbiosis relationship is present in *Dirofilaria* species and not in *A. reconditum* [29]. Although, 75% (9/12) of samples positive only for *D. immitis* DNA were also found to be positive for *Wolbachia* genotype known to be associated with this filarioid, with a significant correlation (Table 3). As previously reported, *Wolbachia* DNA was detected in 64.0% of samples positive for *D. immitis* [29], and in 80% of positive for *D. repens* [48]. In the present study, we investigated the link between *Wolbachia* genotype and *D. repens* infection. Samples positive for a single DNA of *D. repens* had a significant correlation with the *Wolbachia* genotype known to be associated with this filarioid. This result seems to be corroborated by Vytautas et al. [48]. Interestingly, the single DNA of *D. repens* was also strongly correlated with the presence of both *Wolbachia* strains associated with *Dirofilaria* spp. This association could be explained either by the presence of an occult co-infection with *D. immitis* or by an exchange of *Wolbachia* strains between *Dirofilaria* spp. The first suggestion is supported by the fact that co-infection of *D. repens* and *D. immitis* is often associated with an occult form. This phenomenon results from a competitive suppression between microfilariae species [21]. On the other hand, *Wolbachia* of *D. immitis* was detected in 29.63% (8/27) of samples free for *D. immitis* DNA and, in the same samples, an antigen was detected after heat-pre-treatment of sera. This result confirms the possibility to detect *Wolbachia* in occult infection. The utility of *Wolbachia* as a diagnosis target for the occult heartworm was demonstrated in the dead-end host, like human and cat, where the parasite cannot achieve its maturation and the infection might be amicrofilariaemic [47]. However, the second suggestion related to the exchange of *Wolbachia* strains between *Dirofilaria* species contrasts with the published data. The *Wolbachia* transmission principally occurs via eggs of female worms (vertical transmission), unlike the horizontal transfer, which requires a mediation mechanisms, a genetic potential for transfer and suitable environmental conditions [49]. The vertical transfer of *Wolbachia* leads to the specialization of the host-symbiotic relationship [50]. It was experimentally induced by the crossing of *B. malayi* and *B. pahangi* [36]. Theoretically, exchange between *D. immitis* and *D. repens* is hardly possible in natural conditions, because these filariae do not share the same site and the adult worms will not have contact inside the host organism [51]. In addition, it was reported that each genotype of *Wolbachia* has a specific filarial host [37], and the living worms can release their *Wolbachia* into host tissues [39]. We believe that the presence of a determined genotype of *Wolbachia* is a reliable marker for the presence of its filarial host.

### Heartworm antigen detection and performance characteristics assays

In the present study, all diagnosis approaches did not react with samples from the negative control group. We assessed the diagnosis value of ELISA (DiroCHEK®) in detecting heartworm. The direct exploration of heartworm antigen from sera without heating sera showed a moderate performance, where the sensitivity and specificity values do not exceed 65.5% and 93.5%, respectively. Positive antigen tests were obtained from 19 out of 29 (65.52%) of samples determined as true positive for heartworm and often harbored both *D. immitis* and its *Wolbachia* DNA. However, 10 (34.38%) for whom 80% (8) of them harbored only *Wolbachia* of *D. immitis* DNA, remained undetectable by serology. The lack of sensitivity of this assay was unexpected. Nevertheless, similar results have been recently reported, 41 (38.7%) positive PCR-confirmed microfilaremic samples were negative for heartworm antigen [52]. Therefore, *Wolbachia* interact with host by activating the Th1 type protective-immune response [53], which could be implemented in the clearance of heartworm antigens. Finally, 9 out of 28 (32.14%) were positive for *A. reconditum* and/or *D. repens*: these filariae are known to generate a cross reactivity with heartworm antigen test even the absence of heat-treatment of sera [17]. However, 29 (96.67%) samples positive for at least one molecular marker of heartworm that tested positive for heartworm antigen after heating sera, this step has recently been added to improve the sensitivity of this test under certain circumstances [15, 43]. In contrast, the heat-pretreatment of sera alters strongly the specificity of the heartworm antigen test. The cross reactivity was obtained from the overall samples positive for any filarial parasites, as well as samples positive for unknown antigens. Filarial and non-filarial nematodes, such as *Angiostrongylus vasorum*, *Spirocerca lupi*, and *D. repens* and *A. reconditum* are known to be cross react with heartworm antigen tests [18, 55]. Therefore, molecular and serological diagnosis of *A. vasorum* from canine blood showed a close efficiency [56]. Regarding the life cycle of *A. vasorum* [57, 58], the absence of its DNA from all blood samples tested in the present study is not sufficient enough to exonerate the infection. However, the qPCR targeting *A. vasorum* developed in this study can be used as a simplex for detecting parasites larvae from other biological samples, such as stools, pharyngeal swab as well as in intermediate hosts. In addition, the occult heartworm cannot be confirmed by serological tests when presented as a co-infection with another filarial infection that cross-reacts with the test, especially if associated with *D. repens* infection (Fig. 2), this type of case resulting from an unknown mechanism of D. *repens* microfilariae, which shows a suppressive effect on those of *D. immitis* [38]. The ESDA does not recommend the routine heat-pretreatment of sera in area endemic for these parasitosis, as the only recommends it to resolve the discrepancy between the other tests, particularly when the dog is positive on the microfilaria test and negative by serology, and to confirm a suspicion of clinical disease suggestive of microfilaremia [11, 12].

## Conclusion

Herein, we established a multiplexed molecular approach with the potential to detect and identify filariae and their *Wolbachia* DNA from blood samples. We encourage researchers to follow the molecular diagnostic procedure summarized in Figure 3, where the molecular markers of filarial parasites are explored by the pan-filarial and pan-Wolbachia qPCRs, followed by the most common rapid typing method for canine filariasis represented by *A. reconditum*, *D. immitis* and *D. repens* with the specific *Wolbachia* of these latter two filarioids in order to complete the molecular diagnosis. Finally, the classical approach based on sequencing analysis of DNA, is still being considered to identify the etiological agent in the endemic zone for other filarial parasites. The specific detection of *Wolbachia* genotypes could be used for the diagnosis of filariosis and the assessment of related pathogeny within dead-end hosts, as the case of *D. immitis* infection in humans and cats. Furthermore, the quantitative character of the qPCRs offers a suitable tool to assess treatment effectiveness against filarialiosis, as well as against their *Wolbachia*. The use of detection of heartworm antigen without heat treatment is not reliable in an endemic area with other filariasis, including *A. reconditum* and *D. repens*. We discourage the use of heat pretreatment of sera, which significantly alters the specificity of the assay due to the cross-reactivity of many filarial and non-filarial nematodes and the possibility of false-positive results that may induce unnecessary heavy treatment.

## Supporting information

Specificity of Triplex qPCR A: D. immitis DNA, B: D. repens DNA, C: A. reconditum DNA, D: specific and simultaneous detection of three DNA.

Positive control DNA used for sensitivity and specificity determination of designed oligonucleotides.

Negative control DNA used for sensitivity and specificity determination of designed oligonucleotides.

Specificity assessment of Triplex TaqMan qPCR for single and simultaneous detection of D. immitis, D. repens and A. reconditum DNA.

Sensitivity and assay performance characteristics of pan-filarial qPCR system.

Sensitivity and assay performance characteristics of Triplex TaqMan qPCR.

Sensitivity and assay performance of single detection of D. immitis and their Wolbachia using COI-Triplex and ftsZ duplex TaqMan qPCRs, respectively.

Screening followed by sequence typing approach

Distribution of molecular markers

Molecular and serological exploration of canine filariosis in studied groups.

Performance characteristics of molecular and serological approaches for the diagnosis of heartworm.

Performance characteristics of molecular approaches for the diagnosis of D. repens and A. reconditum.

## Acknowledgments

We gratefully thank Sebastien Cortaredona for assistance with statistical analysis.

## Consent for publication

Not applicable.

## Availability of data and material

The data supporting the conclusions of this article are included within the article.

## Authors’ contributions

YL, BD, MV and OM designed the study. YL, BD, OM designed, and YL carried out the data analysis. YL, MV, EAN and OM drafted the manuscript. All authors read and approved the final version of the manuscript.

---

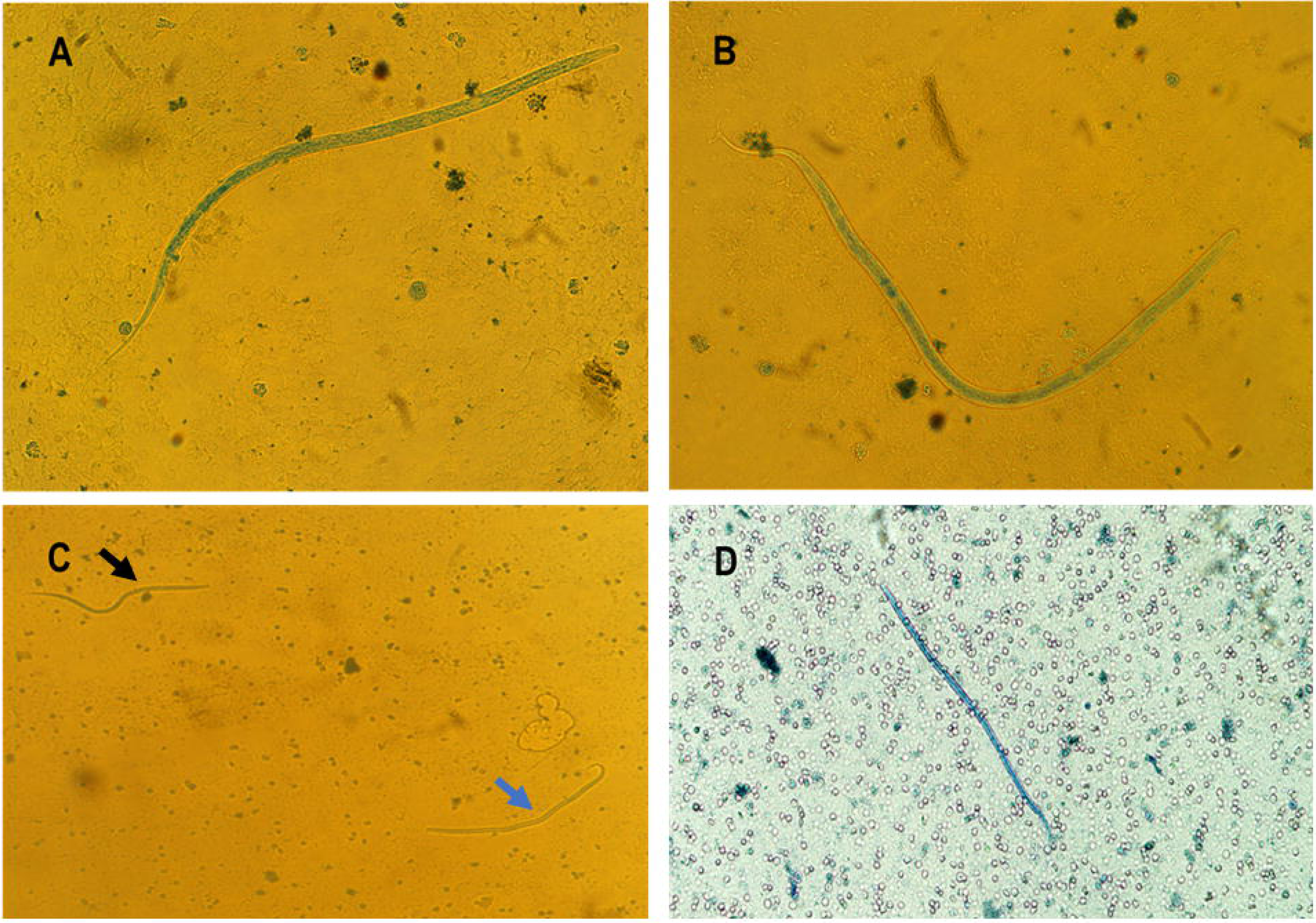

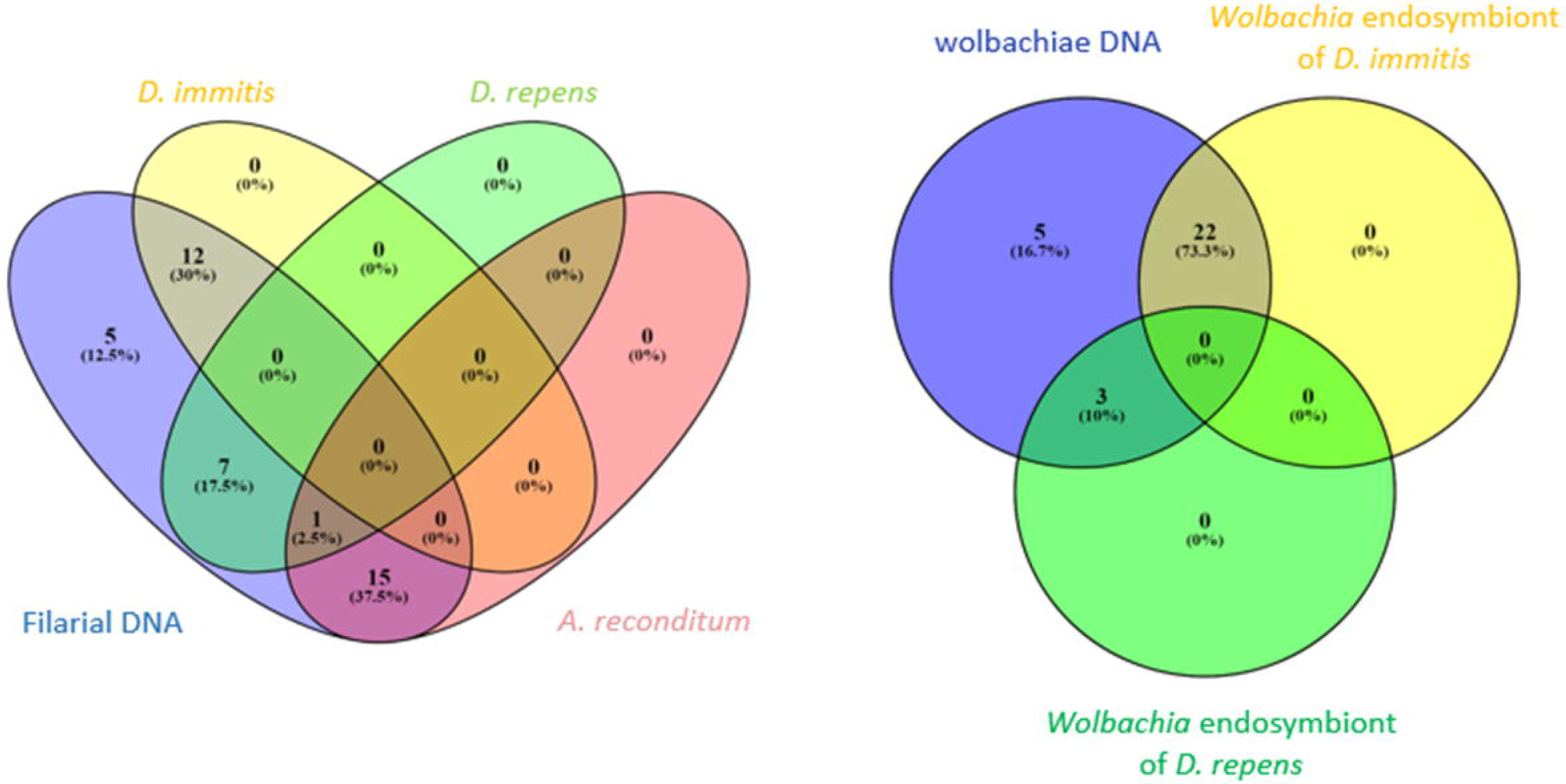

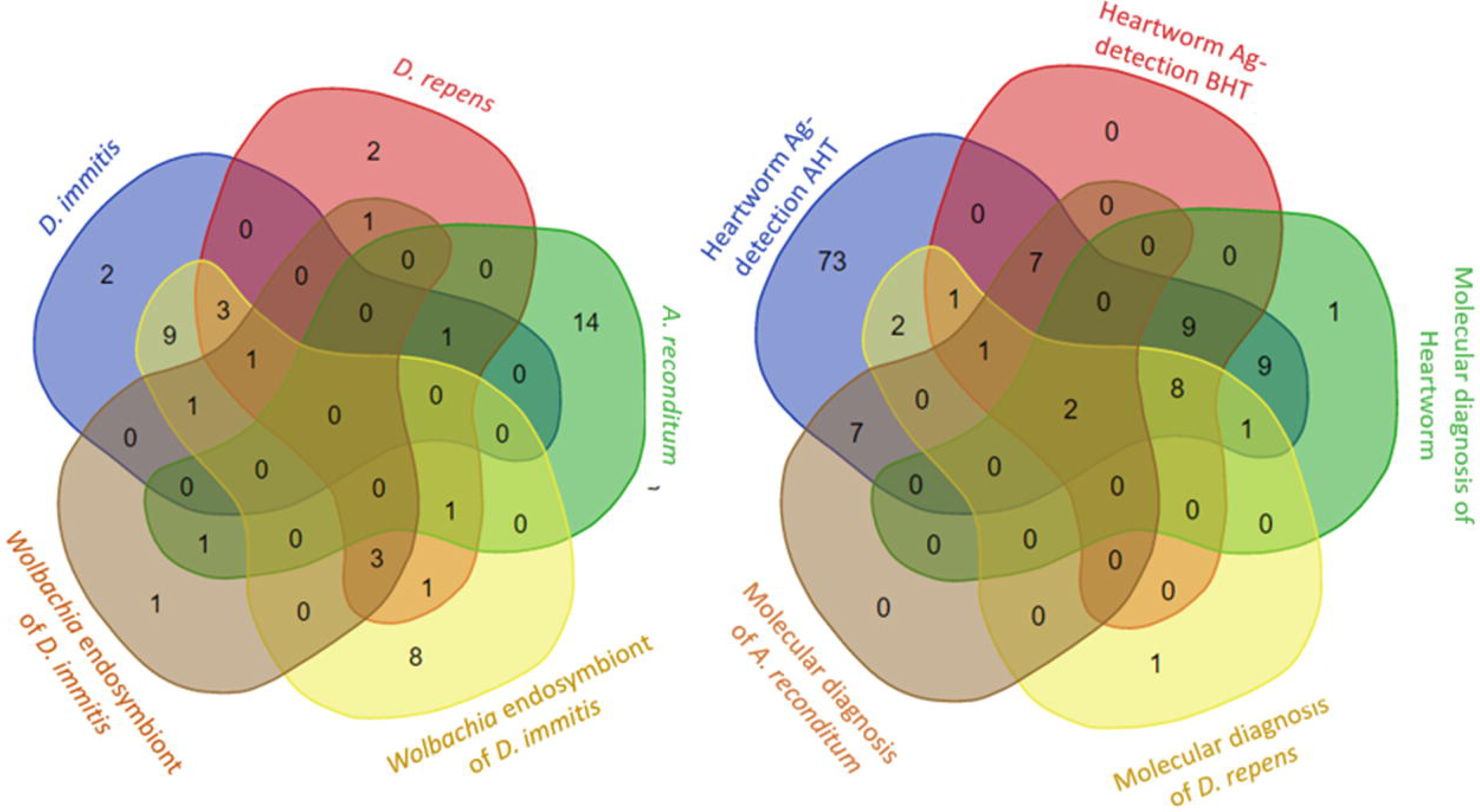

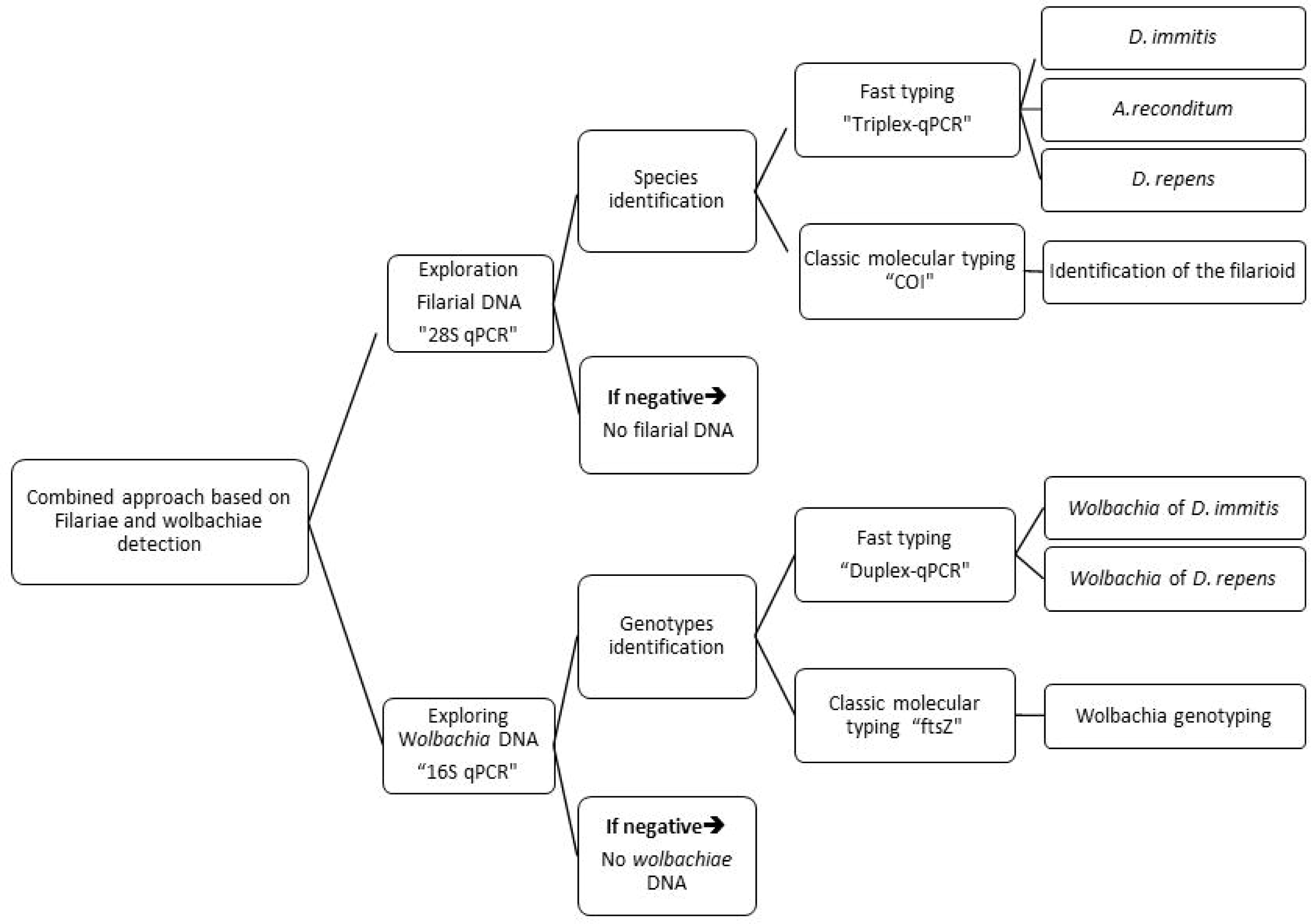

