## Supplementary figures and images for "Development of a multiplexed qPCRs-based approach for the diagnosis of *Dirofilaria immitis, D. repens*, *Acanthocheilonema reconditum* and the others filariosis"

### Specificity of Triplex qPCR A: D. immitis DNA, B: D. repens DNA, C: A. reconditum DNA, D: specific and simultaneous detection of three DNA.

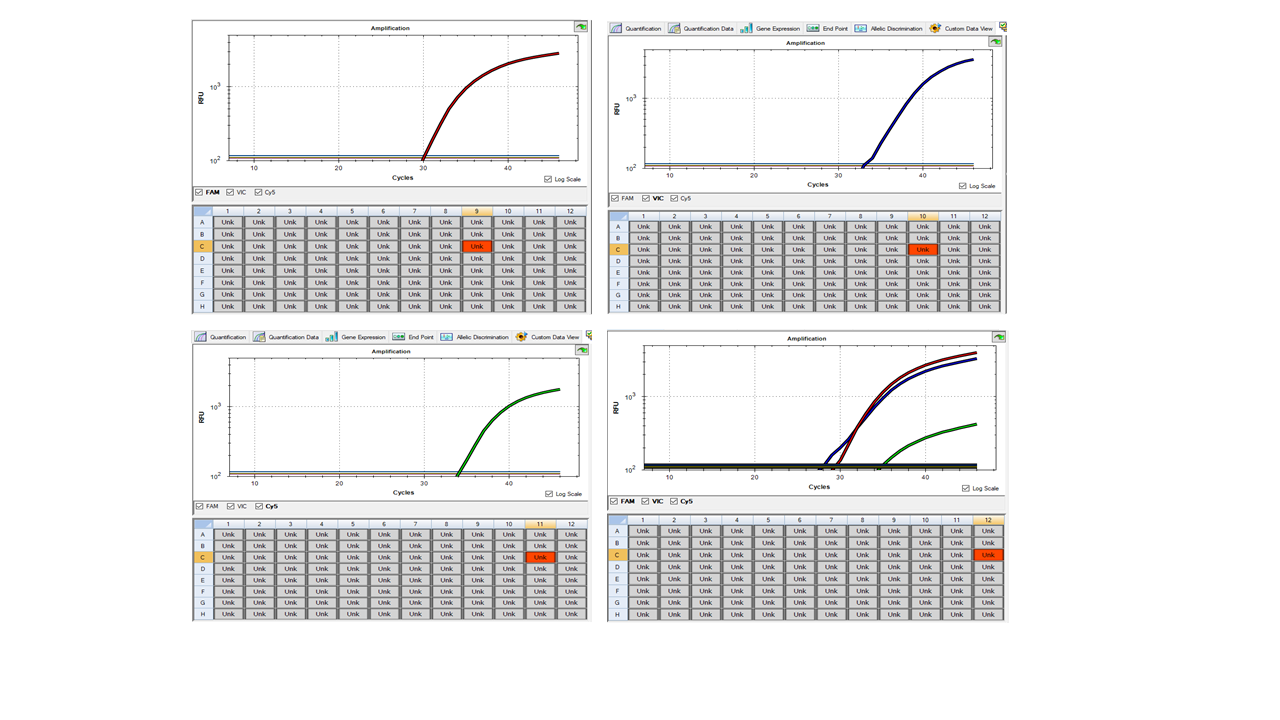

### Supplemental Data 1

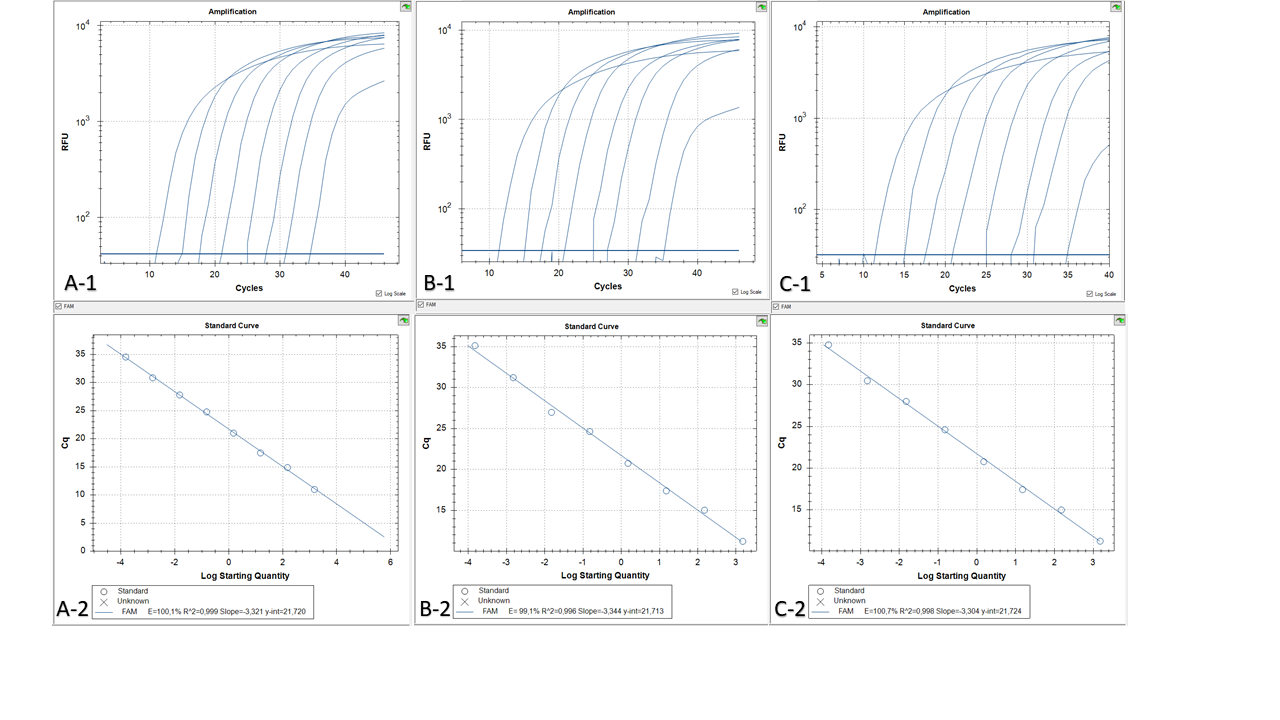

### Supplemental Data 2

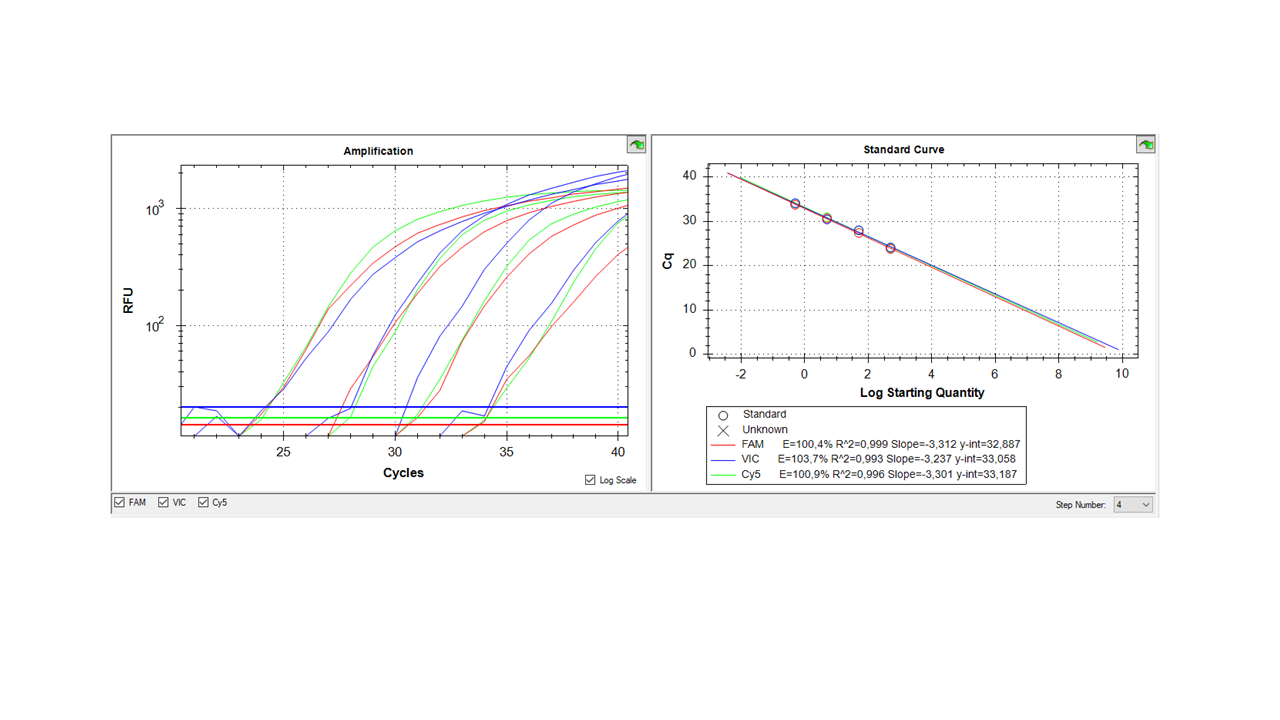

### Supplemental Data 3

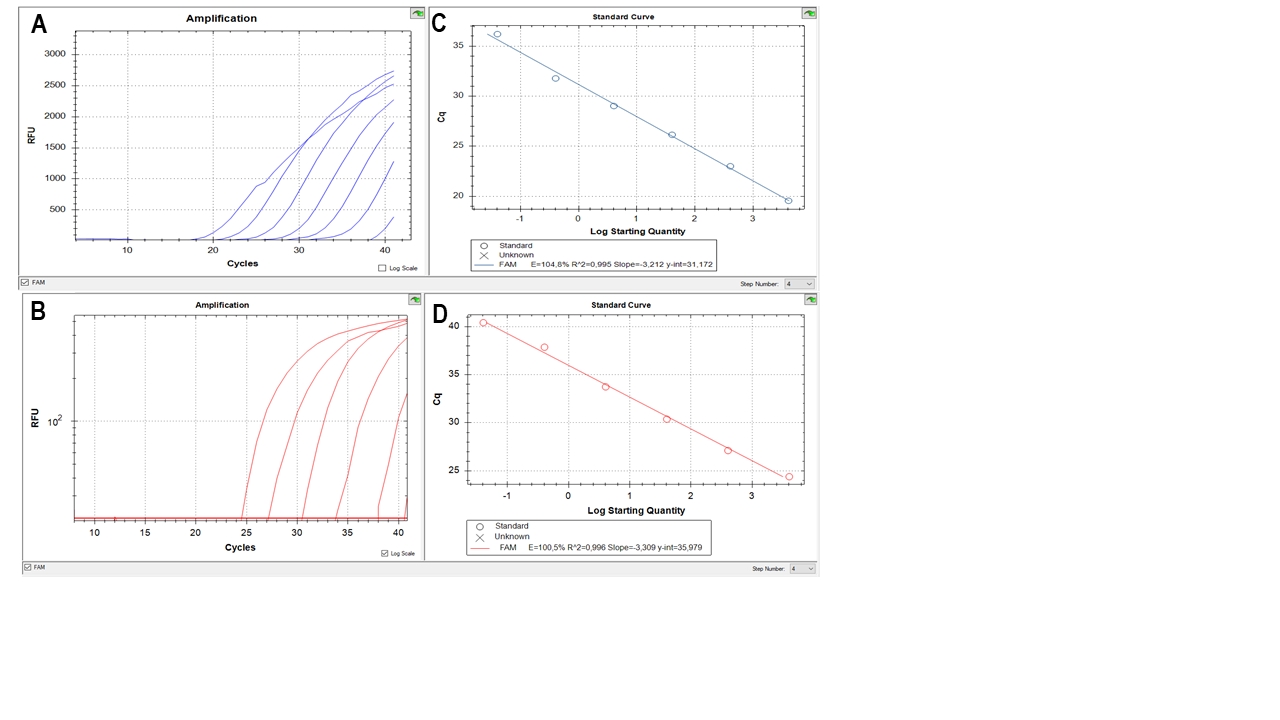
