## Supplementary material for "Development of a multiplexed qPCRs-based approach for the diagnosis of *Dirofilaria immitis, D. repens*, *Acanthocheilonema reconditum* and the others filariosis": Positive control DNA used for sensitivity and specificity determination of designed oligonucleotides.

| N° | **Samples** | **Species name** | **Isolation source** | **Host** | **Country** |
| --- | --- | --- | --- | --- | --- |
| 1 | Adult worm "Deep" | *Dirofilaria immitis* | Adult worm from pulmonary arteries | Dog after necropsy | Marsielle-France |
| 2 | Worm4 | *Dirofilaria repens* | subcutaneous nodules | German Shepherd living in Pau | University Hospital (Dermatology Department, North Hospital, Marseille, France) |
| 3 | Worm1 | *Dirofilaria repens* | Adult worm (peritoneal cavity) | Donkey | Egypt |
| 4 | EA28-EA50…. | *Acanthocheilonema reconditum* | Blood | dog | Cote d'Ivoir |
| 5 | DNA | *Onchocerca lupi* | skin | dog | Italy "Prof Dominico" |
| 6 | DNA | *Thelazia callipaeda* | eye | dog | Italy "Prof Dominico" |
| 7 | MT 91 & MT 90 | *Cercopithifilaria bainae* | Blood | dog | French Guiana |
| 8 | 60 | *Acanthocheilonema sp.* | Blood | Dog | Egypt |
| 9 | 192 | *Dipetalonema Sp.* | Blood | Horse | Egypt |
| 10 | Worm3 | *Dipetalonema Sp.* | Adult worm (peritoneal cavity) | Donkey | Egypt |
| 11 | RC68R | *Spirocerca vulpis* | spleen | Foxes | French |
| 12 | infected Ae. aegypti | *Brugia malayi* | infected Ae. aegypti | Laboratory strain | University of Georgia College of Veterinary Medicine, Athens, (USA). |
| 13 | infected Ae. aegypti | *Brugia pahangi* | infected Ae. aegypti | Laboratory strain | University of Georgia College of Veterinary Medicine, Athens, (USA). |
| 14 | Monkey blood B6 | *Brugia sp.* | Blood | Howler monkeys | French Guiana |
| 15 | Human filariasis "A" | *Wuchereria bancrofti* | blood | Human | Cote d'Ivoir |
| 16 | Human filariasis "B" | *Loa loa* | Blood | Gabonese children | Gabon |
| 17 | Human filariasis "C" | *Mansonella perstens* | Blood | febrile patient | Senegal |
| 18 | Monkey blood B8 | *Onchocerca cervicalis* | Blood | Howler monkeys | French Guiana |
| 19 | Worm1 | *Setaria digitata* | Adult worm (peritoneal cavity) | Donkey | Egypt |
| 20 | 32/44/49/66/67/78/79/83/93 | *Dipetalonema Sp.* | Blood | Donkey | Egypt |
| 21 | Adult worm "CH11 & CH31" | *Abreviata caucasica* | stool of chimpanzee | Wild chimpanzees | Senegal |
| 22 | worm | *Ascaris sp.* | adult worm | wild foxes | French |
| 23 | worm | *Toxocara catis* | adult worm | cat | French |
| 24 | micro-worm | *Pangrellus sp.* | micro-worm | culture | IHU |
