## Supplementary material for "Development of a multiplexed qPCRs-based approach for the diagnosis of *Dirofilaria immitis, D. repens*, *Acanthocheilonema reconditum* and the others filariosis": Negative control DNA used for sensitivity and specificity determination of designed oligonucleotides.

| **N°** | **Organism** | **Name** |
| --- | --- | --- |
| 1 | Protozoa | *Trypanosoma evansi** |
| 2 | Protozoa | *Trypanosoma brucei brucei** |
| 3 | Protozoa | *Trypanosoma brucei gambiense* |
| 4 | Protozoa | *Trypanosoma brucei gambiense* biyiamina groupe II*** |
| 5 | Protozoa | *Trypanosoma congolense* IL 3000*** |
| 7 | Protozoa | *Trypanosoma vivax ** |
| 8 | Protozoa | *Trypanosoma cruzi CL Brunner** |
| 9 | Protozoa | *Trypanosoma cruzi (Dog KIM)** |
| 10 | Protozoa | *Leptomonas* spp. *** |
| 12 | Bacteria | *Rickettsia montanensis* |
| 14 | Lice | Head lice *(Pediculus humanus capitis)* |
| 15 | Bacteria | *Ehrlichia canis* |
| 16 | Bacteria | *Coxiella burnetii* |
| 17 | Bacteria | *Borrelia recurrentis* |
| 18 | Bacteria | *Staphylococcus hominis* |
| 19 | Bacteria | *Asaia bogorensis* |
| 20 | Piroplasm | *Babesia canis (*Moocky*)* |
| 21 | Bacteria | *Haemophilus influenzae* |
| 22 | Bacteria | *Asaia* C2 |
| 23 | Bacteria | *Anaplasma phagocytophilum* |
| 24 | Bacteria | *Enterobacter aerogenes* |
| 25 | Bacteria | *Streptococcus pneumoniae* |
| 27 | Bacteria | *Salmonella enterica* |
| 29 | Bacteria | *Citrobacter koseri* |
| 30 | Bacteria | *Gardnerella vaginalis* |
| 31 | Bacteria | *Streptococcus pyogenes* |
| 32 | Protozoa | *Plasmodium* sp. |
| 33 | Bacteria | *Rickettsia typhi* |
| 34 | Bacteria | *Enterococcus faecium* |
| 35 | Bacteria | *Streptococcus agalactiae* |
| 36 | Bacteria | *Rickettsia conorii* |
| 37 | Tick | *Rhipicephalus microplus* |
| 38 | Bovine | BA286 cell line |
| 39 | Dog | DH62 cell line |
| 40 | Human | HL60 cell line *(Homo sapiens)* |
| 41 | Donkey | ANE 4 *(Equus asinus)* |
| 42 | Horse | CV.G 22 *(Equs caballus)* |
| 43 | Flea | *Ctenocephalides felis* |
| 44 | Bedbugs | *Cimex lectularius* |
| 45 | Tick | *Hyalomma marginatum* |
| 46 | Tick | *Amblyomma variegatum* |
| 47 | Dog | DH62 dog cell line (*Canis lupus familiaris*) |
| 48 | Protozoa | *Hepatozoon canis* |
| 49 | Protozoa | *Leptomonas saimouri* |
| 50 | Piroplasm | *Theileria equi* |
| 51 | Bacteria | *Rickettsia massiliae* |
| 52 | Bacteria | *Staphylococcus haemolyticus* |
| 53 | Bacteria | *Staphylococcus aureus* |
| 54 | Bacteria | *Rickettsia felis* |
| 55 | Bacteria | *Stenotrophomonas maltophilia* |
| 56 | Bacteria | *Acinetobacter sp.* |
| 57 | Bacteria | *Enterobacter aerogenes* |
