## Supplementary material for "Development of a multiplexed qPCRs-based approach for the diagnosis of *Dirofilaria immitis, D. repens*, *Acanthocheilonema reconditum* and the others filariosis": Specificity assessment of Triplex TaqMan qPCR for single and simultaneous detection of D. immitis, D. repens and A. reconditum DNA.

|  | FAM  End RFUs | VIC  End RFUs | Cy5  End RFUs | Positive signal |
| --- | --- | --- | --- | --- |
| *D.immitis* DNA | ***2619** | 49.6 | 5.13 | FAM |
| *D.repens* DNA | 71 | ***3085** | 9.23 | VIC |
| *A. reconditum* DNA | 19.5 | 56,1 | ***1583** | Cy5 |
| Pooled DNA | ***3603** | ***2973** | ***375** | FAM-VIC-Cy5 |
| Cut Off Value | 351.3 | 357.3 | 99.2 | // |
| Negative control | 35,7 | 49,1 | 0,589 | // |

**RFU**: relative fluorescence unit; *: positive reaction (greater than the threshold).

**RFU**: relative fluorescence unit; *: positive reaction (greater than the threshold).
