## Supplementary material for "Development of a multiplexed qPCRs-based approach for the diagnosis of *Dirofilaria immitis, D. repens*, *Acanthocheilonema reconditum* and the others filariosis": Sensitivity and assay performance characteristics of pan-filarial qPCR system.

| **Mf concentration** | | | | ***D. immitis DNA*** | | | | | | ***D. repens DNA*** | | | | | | ***A. reconditum DNA*** | | | | |
| --- | --- | --- | --- | --- | --- | --- | --- | --- | --- | --- | --- | --- | --- | --- | --- | --- | --- | --- | --- | --- |
|  |  |  |  |  | **Ct** | **E-RFU** |  | **SCRS** |  |  | **Cq** | **E-RFU** |  | **SCRS** |  |  | **Cq** | **E-RFU** |  | **SCRS** |
|  | **DNA from mfs/5µl** | **mfs/ml** |  |  |  |  |  |  |  |  |  |  |  |  |  |  |  |  |  |  |
|  | **0.75E+01** | **1.5 E+03** |  |  | **12.32** | **6127** |  | **(E= 104.3%)**  **(S=-3.22)**  **(Y.int;= 23.15)**  **(R^2=0.997)** |  |  | **11.26** | **5938** |  | **E=99.1%)**  **(S=-3.34)**  **(Y.int.=21.71)**  **(R^2=0.996)** |  |  | **11.26** | **5667** |  | **(E=100.7%)**  **(S=-3.304)**  **(Y.int.=21.72)**  **(R^2=0.998)** |
|  | **0.75E+00** | **1.5 E+02** |  |  | **16.1** | **7420** |  |  |  |  | **15** | **7802** |  |  |  |  | **15** | **7619** |  |  |
|  | **0.75E-01** | **1.5 E+01** |  |  | **19.44** | **7690** |  |  |  |  | **17.44** | **8399** |  |  |  |  | **17.44** | **7825** |  |  |
|  | **0.75E-02** | **1.5 E+00** |  |  | **22.82** | **8221** |  |  |  |  | **20.78** | **9037** |  |  |  |  | **20.78** | **8270** |  |  |
|  | **0.75E-03** | **1.5 E-01** |  |  | **26.12** | **7660** |  |  |  |  | **24.6** | **7623** |  |  |  |  | **24.6** | **7794** |  |  |
|  | **0.75E-04** | **1.5 E-02** |  |  | **29.58** | **7024** |  |  |  |  | **28** | **7389** |  |  |  |  | **28** | **6402** |  |  |
|  | **0.75E-05** | **1.5 E-03** |  |  | **32.23** | **5360** |  |  |  |  | **30.48** | **5595** |  |  |  |  | **30.48** | **5435** |  |  |
|  | **0.75E-06** | **1.5 E-04** |  |  | **34.66** | **2325** |  |  |  |  | **34.78** | **1226** |  |  |  |  | **34.78** | **2625** |  |  |
|  | **Cut Off Value** | **<1.5 E-04** |  |  | **35** | **909** |  |  |  |  | **35** | **909** |  |  |  |  | **35** | **909** |  |  |
|  | **Negative Control** | **0** |  |  | **N/A** | **10,1** |  |  |  |  | **N/A** | **10,1** |  |  |  |  | **N/A** | **10,1** |  |  |

**mfs**: microfilariae, **Cq**: cycle quantification value; **N/A**: No amplification, **E-RFU**: End of relative fluorescence unit, **SCRS**: Standard Curve Results Spreadsheet, **E**: Efficiency, **S**: Slope, **Y.int:** Y-intercept.
