## Supplementary material for "Development of a multiplexed qPCRs-based approach for the diagnosis of *Dirofilaria immitis, D. repens*, *Acanthocheilonema reconditum* and the others filariosis": Sensitivity and assay performance characteristics of Triplex TaqMan qPCR.

| Mf concentration | | | | *D. immitis DNA* | | | | | | *D. repens DNA* | | | | | | *A. reconditum DNA* | | | | |
| --- | --- | --- | --- | --- | --- | --- | --- | --- | --- | --- | --- | --- | --- | --- | --- | --- | --- | --- | --- | --- |
|  |  |  |  |  | Cq | End RFU FAM |  | SCRS |  |  | Cq | End RFU VIC |  | SCRS |  |  | Cq | End RFU Cy-5 |  | SCRS |
|  | DNA from mfs/5µl | mfs/ml of each species |  |  |  |  |  |  |  |  |  |  |  |  |  |  |  |  |  |  |
|  | 2.5E+00 | 5.00E+02 |  |  | 23,81 | 1387 |  | (E= 100.4%)  (S=-3.312)  (Y.int= 32.89)  (R^2=0.999) |  |  | 24,13 | 1590 |  | (E=103.7%)  (S=-3.236)  (Y.int=33.06)  (R^2=0.993) |  |  | 24,03 | 1405 |  | E=100.9%)  (S=-3.30)  (Y.int=33.212)  (R^2=0.996) |
|  | 2.5E-01 | 5.00E+01 |  |  | 27,41 | 1241 |  |  |  |  | 28,01 | 1852 |  |  |  |  | 27,94 | 1302 |  |  |
|  | 2.5E-02 | 5.00E+00 |  |  | 30,7 | 865 |  |  |  |  | 30,45 | 1614 |  |  |  |  | 30,9 | 1009 |  |  |
|  | 2.5E-03 | 5.00E-01 |  |  | 33,75 | 297 |  |  |  |  | 34,11 | 567 |  |  |  |  | 34,05 | 512 |  |  |
|  | 2.5E-04 | 5.00E-02 |  |  | 35,83 | 193 |  |  |  |  | 36,45 | 66,1 |  |  |  |  | 38,74 | 11,3 |  |  |
|  | Cut Off Value | // |  |  | 35 | 200 |  |  |  |  | 35 | 73.9 |  |  |  |  | 35 | 40.2 |  |  |
|  | Negative Control | // |  |  | // | 10,5 |  |  |  |  | // | 15 |  |  |  |  | // | 1.5 |  |  |

**mfs**: microfilariae, **Cq**: cycle quantification value, **N/A**: No amplification, **RFU**: relative fluorescence unit, **SCRS**: Standard Curve Results Spreadsheet, **E**: Efficiency, **S**: Slope, **Y.int:** Y-intercept.
