## Supplementary material for "Development of a multiplexed qPCRs-based approach for the diagnosis of *Dirofilaria immitis, D. repens*, *Acanthocheilonema reconditum* and the others filariosis": Sensitivity and assay performance of single detection of D. immitis and their Wolbachia using COI-Triplex and ftsZ duplex TaqMan qPCRs, respectively.

|  | | | | Single detection of *D. immitis DNA using* triplex qPCR system | | | | | Single detection of DNA of *Wolbachia* of *D. immitis DNA using* duplex qPCR system | | | |
| --- | --- | --- | --- | --- | --- | --- | --- | --- | --- | --- | --- | --- |
|  | DNA from  mfs/5µl | mfs/ml of each species |  |  | Ct | End RFU | SCRS |  |  | Mct | End RFU | SCRS |
|  | Mother solution | 4,03E+03 |  |  | *18.11 | 2383 | (E=104.8%)  (S=-3.212)  (Y.int.= 31.172)  (R^2=0.995) |  |  | *24.39 | 497 | (E=100.5%)  (S=-3.3)  (Y.int.=35.98)  (R^2=0.996) |
|  | 2.01E+01 | 4,03E+02 |  |  | *21.03 | 2591 |  |  |  | *27.11 | 450 |  |
|  | 2.01E+00 | 4,03E+01 |  |  | *24.66 | 2448 |  |  |  | *30.38 | 455 |  |
|  | 2.01E-01 | 4,03E+00 |  |  | *27.54 | 2006 |  |  |  | *33.74 | 272 |  |
|  | 2.01E-02 | 4,03E-01 |  |  | *29.23 | 1504 |  |  |  | *37.87 | 67 |  |
|  | 2.01E-03 | 4,03E-02 |  |  | *30.93 | 773 |  |  |  | 40.42 | 14.9 |  |
|  | 2.01E-04 | 4,03E-03 |  |  | 35.28 | 125 |  |  |  | N/A | N/A |  |
|  | 2.01E-05 | 4,03E-04 |  |  | 38.39 | 21 |  |  |  | N/A | N/A |  |
|  | Cut Off value | // |  |  | 35.5 | 158 |  |  |  | 38 | 30.5 |  |
|  | Negative Control | // |  |  | // | 8.68 |  |  |  | // | 5 |  |

**N/A**: No amplification, **RFU**: relative fluorescence unit, **SCRS**: Standard Curve Results Spreadsheet, **E**: Efficiency, **S**: Slope, **Y.int:** Y-intercept, *****: positive reaction.
