## Supplementary material for "Development of a multiplexed qPCRs-based approach for the diagnosis of *Dirofilaria immitis, D. repens*, *Acanthocheilonema reconditum* and the others filariosis": Screening followed by sequence typing approach

| **Names** | **Total** | **Sample code** |
| --- | --- | --- |
| - A. reconditum COI sequences | 1 | mf-CI-122 |
| - D. repens COI sequences |  |  |
| - Wolbachia of D. immitis ftsZ sequences |  |  |
| - D. immitis COI sequences | 9 | CI-EA-136, mf-MWD19, CI-EA-26, mf-MWD03, CI-EA-135, CI-EA-01, CI-EA-03, CI-EA-31, CI-EA-132 |
| - Wolbachia of D. immitis ftsZ sequences |  |  |
| - D. immitis COI sequences | 1 | CI-EA-64 |
| - Untyped Wolbachiae DNA |  |  |
| - D. repens COI sequences | 1 | CI-EA-120 |
| - Wolbachia of D. immitis ftsZ sequences |  |  |
| - D. repens COI sequences | 1 | CI-EA-56 |
| - Wolbachia of D. repens ftsZ sequences |  |  |
| - D. repens COI sequences | 3 | mf-MWD05, mf-MWD02, mf-MWD09 |
| - Untyped Wolbachiae DNA |  |  |
| - A. reconditum COI sequences | 1 | mf-CI-97 |
| - Wolbachia of D. repens ftsZ sequences |  |  |
