## Supplementary material for "Development of a multiplexed qPCRs-based approach for the diagnosis of *Dirofilaria immitis, D. repens*, *Acanthocheilonema reconditum* and the others filariosis": Distribution of molecular markers

| **Markers** | **total detected** | **Sample names** |
| --- | --- | --- |
| - *D. immitis* - *D. repens* - *Wolbachia of D. immitis* - *Wolbachia of D. repens* | 1 | mf-MWD01 |
| - *A. reconditum* - *D. immitis* - *D. repens* | 1 | mf-CI-123 |
| - *D. immitis* - *D. repens* - *Wolbachia of D. immitis* | 3 | CI-EA-122 CI-EA-91 CI-EA-55 |
| - *D. immitis* - *Wolbachia of D. immitis* - *Wolbachia of D. repens* | 1 | CI-EA-64 |
| - *A. reconditum* - *D. repens* - *Wolbachia of D. immitis* | 1 | mf-CI-122 |
| - *D. repens* - *Wolbachia of D. immitis* - *Wolbachia of D. repens* | 3 | mf-MWD05, mf-MWD02, mf-MWD09 |
| - *D. immitis* - *Wolbachia of D. immitis* | 9 | CI-EA-136, mf-MWD19, CI-EA-26, mf-MWD03, CI-EA-135, CI-EA-01, CI-EA-03, CI-EA-132, CI-EA-31 |
| - *D. repens* - *Wolbachia of D. immitis* | 1 | CI-EA-120 |
| - *D. repens* - *Wolbachia of D. repens* | 1 | CI-EA-56 |
| - *A. reconditum* - *Wolbachia of D. repens* | 1 | mf-CI-97 |
| - *D. immitis* | 2 | CI-EA-97, CI-EA-27 |
| - *D. repens* | 2 | CI-EA-62, CI-EA-124 |
| - *A. reconditum* | 14 | mf-CI-50, mf-CI-109, mf-CI-28, mf-CI-118, mf-CI-101, mf-CI-110, mf-CI-115, mf-CI-104, mf-CI-111, mf-CI-116, mf-CI-102, mf-CI-98, mf-CI-99, mf-CI-96 |
| - *Wolbachia of D. immitis* | 8 | CI-EA-121, CI-EA-111, CI-EA-119, CI-EA-116, CI-EA-113, CI-EA-94, CI-EA-117, CI-EA-114 |
| - *Wolbachia of D. repens* | 1 | MWD09 |
