## Supplementary material for "Development of a multiplexed qPCRs-based approach for the diagnosis of *Dirofilaria immitis, D. repens*, *Acanthocheilonema reconditum* and the others filariosis": Molecular and serological exploration of canine filariosis in studied groups.

| **Markers** | **total detected** | **Sample names** |
| --- | --- | --- |
| - A. reconditum infection - Ag-detection AHT - Ag-detection BHT - D. repens infection - Heartworm infection | 2 | mf-CI-122, mf-CI-123 |
| - Ag-detection AHT - Ag-detection BHT - D. repens infection - Heartworm infection | 8 | CI-EA-122, mf-MWD02, mf-MWD09, CI-EA-64, CI-EA-91, mf-MWD01, CI-EA-55, CI-EA-120 |
| - A. reconditum infection - Ag-detection AHT - Ag-detection BHT - D. repens infection | 1 | mf-CI-97 |
| - Ag-detection AHT - Ag-detection BHT - Heartworm infection | 9 | mf-MWD19, CI-EA-26, mf-MWD03, CI-EA-135, CI-EA-01, CI-EA-132, CI-EA-31, CI-EA-97, CI-EA-136 |
| - Ag-detection AHT - Ag-detection BHT - D. repens infection | 1 | CI-EA-62 |
| - A. reconditum infection - Ag-detection AHT - Ag-detection BHT | 7 | mf-CI-118, mf-CI-104, mf-CI-111, mf-CI-116, mf-CI-99, mf-CI-101, mf-CI-115 |
| - Ag-detection AHT - D. repens infection - Heartworm infection | 1 | mf-MWD05 |
| - Ag-detection AHT - Heartworm infection | 9 | CI-EA-121, CI-EA-111, CI-EA-119, CI-EA-116, CI-EA-117, CI-EA-114, CI-EA-94, CI-EA-03, CI-EA-27 |
| - Ag-detection AHT - D. repens infection | 2 | CI-EA-56, CI-EA-124 |
| - A. reconditum infection - Ag-detection AHT | 7 | mf-CI-109, mf-CI-28, mf-CI-102, mf-CI-50, mf-CI-110, mf-CI-98, mf-CI-96 |
| - Ag-detection AHT | 73 | CI-EA-25, CI-EA-82, CI-EA-102, CI-EA-02, CI-EA-28, CI-EA-95, CI-EA-29, CI-EA-21, CI-EA-09, CI-EA-52, CI-EA-133, CI-EA-70, CI-EA-123, CI-EA-104, CI-EA-134, CI-EA-17, CI-EA-118, CI-EA-04, CI-EA-88, CI-EA-67, CI-EA-38, CI-EA-129, CI-EA-84, CI-EA-108, CI-EA-39, CI-EA-96, CI-EA-42, CI-EA-106, CI-EA-103, CI-EA-87, CI-EA-66, CI-EA-131, CI-EA-83, CI-EA-05, CI-EA-22, CI-EA-59, CI-EA-128, CI-EA-99, CI-EA-107, CI-EA-43, CI-EA-100, CI-EA-32, CI-EA-112, CI-EA-13, CI-EA-74, CI-EA-101, CI-EA-46, CI-EA-44, CI-EA-109, CI-EA-54, CI-EA-45, CI-EA-79, CI-EA-90, CI-EA-130, CI-EA-10, CI-EA-98, CI-EA-30, CI-EA-92, CI-EA-110, CI-EA-06, CI-EA-35, CI-EA-41, CI-EA-40, CI-EA-126, CI-EA-53, CI-EA-78, CI-EA-23, CI-EA-37, CI-EA-36, CI-EA-115, CI-EA-125, CI-EA-93, CI-EA-127 |
| - Heartworm infection | 1 | CI-EA-113 |
| - D. repens infection | 1 | Ms-MWD09 |
