## Supplementary material for "Development of a multiplexed qPCRs-based approach for the diagnosis of *Dirofilaria immitis, D. repens*, *Acanthocheilonema reconditum* and the others filariosis": Performance characteristics of molecular and serological approaches for the diagnosis of heartworm.

|  | Molecular approachs | | Heartworm antigen detection | |
| --- | --- | --- | --- | --- |
|  | Sequence typing | Multiplexing qPCRs | B h-Tx | A h-Tx |
| Statistic | Value (%) | Value (%) | Value (%) | Value (%) |
| Correct classification | 96,4 +/- (5,6) | 99,4 +/- (0,9) | 88,7 +/- (4,8) | 45,8 +/- (7,5) |
| Misclassification | 3,6 +/- (5,6) | 0,6 +/- (0,9) | 11,3 +/- (4,8) | 54,2 +/- (7,5) |
| Sensitivity | 82,8 +/- (29,9) | 100,0 +/- (6,0) | 65,5 +/- (18,2) | 100,0 +/- (6,0) |
| Specificity | 99,3 +/- (3,9) | 99,3 +/- (2,0) | 93,5 +/- (4,5) | 34,5 +/- (8,2) |
| False positive rate | 0,7 +/- (2,1) | 0,7 +/- (1,1) | 6,5 +/- (4,1) | 65,5 +/- (7,9) |
| False negative rate | 17,2 +/- (27,5) | 0,0 +/- (0,0) | 34,5 +/- (17,3) | 0,0 +/- (0,0) |
| True prevalence | 17,3 +/- (11,4) | 17,3 +/- (5,7) | 17,3 +/- (5,7) | 17,3 +/- (5,7) |
| Apparent prevalence | 14.88 | 17.85 ± () | 16.67 | 71.43 |
| PPV (Positive Predictive Value) | 96,0 +/- (11,7) | 96,7 +/- (4,9) | 67,9 +/- (17,3) | 24,2 +/- (7,7) |
| NPV (Negative Predictive Value) | 96,5 +/- (6,0) | 100 +/- (0,0) | 92,9 +/- (4,3) | 100,0 +/- (0,0) |
| Cohen's kappa | 0,87 | 0,98 | 0,60 | 0,15 |

**B h-Tx**: Before heat-treatment; **A h-Tx**: After heat-treatment*.*
