## Supplementary material for "Development of a multiplexed qPCRs-based approach for the diagnosis of *Dirofilaria immitis, D. repens*, *Acanthocheilonema reconditum* and the others filariosis": Performance characteristics of molecular approaches for the diagnosis of D. repens and A. reconditum.

|  | *D. repens* | | *A. reconditum* | |
| --- | --- | --- | --- | --- |
|  | Sequence typing | Multiplexing qPCRs | sequencing | Multiplexing |
| Statistic | Value (%) | Value (%) | Value (%) | Value (%) |
| Correct classification | 96,4 +/- (5,6) | 100,0 +/- (0,0) | 99,4 +/- (0,9) | 100,0 +/- (0,0) |
| Misclassification | 3,6 +/- (5,6) | 0,0 +/- (0,0) | 0,6 +/- (0,9) | 0,0 +/- (0,0) |
| Sensitivity | 62,5 +/- (49,4) | 100,0 +/- (10,3) | 94,1 +/- (14,3) | 100,0 +/- (9,8) |
| Specificity | 100,0 +/- (2,4) | 100,0 +/- (1,2) | 100,0 +/- (1,2) | 100,0 +/- (1,2) |
| False positive rate | 0,0 +/- (0,0) | 0,0 +/- (0,0) | 0,0 +/- (0,0) | 0,0 +/- (0,0) |
| False negative rate | 37,5 +/- (47,4) | 0,0 +/- (0,0) | 5,9 +/- (8,5) | 0,0 +/- (0,0) |
| Prevalence | 9,5 +/- (8,9) | 9,5 +/- (4,4) | 10,1 +/- (4,6) | 10,1 +/- (4,6) |
| PPV (Positive Predictive Value) | 100,0 +/- (0,0) | 100,0 +/- (0,0) | 100,0 +/- (0,0) | 100,0 +/- (0,0) |
| NPV (Negative Predictive Value) | 96,2 +/- (6,0) | 100,0 +/- (0,0) | 99,3 +/- (1,0) | 100,0 +/- (0,0) |
| Cohen's kappa | 0,75 | 1,00 | 0,97 | 1,00 |

**B h-Tx**: Before heat-treatment; **A h-Tx**: After heat-treatment*.*
